# Charting the Normal Development of Structural Brain Connectivity in Utero using Diffusion MRI

**DOI:** 10.1101/2025.09.14.676101

**Authors:** Davood Karimi, Bo Li, Athena Taymourtash, Camilo Jaimes, Ellen P. Grant, Simon K. Warfield

## Abstract

Understanding the structural connectivity of the human brain during fetal life is critical for uncovering the early foundations of neural function and vulnerability to developmental disorders. Diffusion-weighted MRI (dMRI) enables non-invasive mapping of white matter pathways and construction of the brain’s structural connectome, but its application to the fetal brain has been limited by data scarcity and technical difficulties in analyzing fetal dMRI data. Here, we present the largest study to date of in utero brain connectivity, analyzing high-quality dMRI data from 198 fetuses between 22 and 37 gestational weeks from the Developing Human Connectome Project. We employed advanced fetal-specific tools for brain segmentation and parcellation, and used ensemble tractography to encourage more complete reconstruction of various white matter tracts. For connection weighting, we relied on the notion of fiber bundle capacity. We reconstructed individual structural connectomes and characterized the developmental trajectories. Graph-theoretical analysis revealed consistent increases in integration and segregation metrics over gestation, while bootstrapping confirmed the robustness of nodal and edgewise developmental patterns. Furthermore, we proposed a novel method for constructing age–specific connectome templates based on aggregation of individual subject connectomes. The new method follows an optimization-based approach to ensure that the connectome templates closely represent individual subject connectomes, are temporally consistent, and proportionally preserve short and long connections. Our analysis shows that this approach is superior to spatial alignment and averaging of the data in image space, with the resulting connectome templates supporting accurate prediction of the gestational age of individual fetuses (mean error= 0.90 ± 0.81 weeks). In summary, this study provides robust normative benchmarks for fetal brain structural connectivity and demonstrates that connectome-based measures capture meaningful developmental signatures, offering a framework for future studies of early brain development.

## 1 INTRODUCTION

Understanding the brain’s structural connectivity is a fundamental goal in neuroscience [1, 2, 3, 4]. The structural connectome underpins and supports all brain activities. Moreover, it can influence and be influenced by the brain’s response to diseases [5]. Diffusion-weighted MRI (dMRI) is a unique non-invasive tool for studying the brain’s structural connectivity [6, 7, 8, 9]. The dMRI signal can be used to estimate the local orientation of white matter fibers in the brain. Building on this local information, tractography algorithms can be applied to trace virtual streamlines connecting predefined brain regions. These streamlines, typically weighted based on measures of tissue microstructure integrity or fiber bundle capacity, determine the strength of connections between the regions to compute a structural connectome [10, 11, 12]. The connectome may be regarded as a mathematical *graph*, where regions represent the graph nodes and the connections represent the edges. Graph-theoretic metrics can subsequently be computed to quantitatively analyze the connectome [13, 4, 7]. A growing body of studies have shown that structural connectivity plays a central role in normal brain development, aging, and degeneration [14, 11, 15, 16, 17, 18]. Although dMRI-based connectivity analysis faces several technical challenges, methodological advances have improved its accuracy and reproducibility [19, 12, 20]. Moreover, our understanding of the capabilities and limitations of this method has substantially improved, enabling more reliable interpretation of the results [21, 22, 23, 24].

However, prior works on structural connectivity have almost exclusively focused on postnatal and adult brains. By comparison, the fetal brain has received far less attention [25, 26, 27, 28, 29, 30, 31, 32, 33]. This has been, in part, due to the limited availability of fetal dMRI data. While there have been large publicly available datasets of pediatric and adult brain dMRI, until recently similar datasets for fetal brain have been lacking. Fetal dMRI also suffers from lower signal to noise ratio, unpredictable motion, and typically shorter scan times [8, 34, 35]. Moreover, due to the incomplete myelination and relatively higher water content, the fetal brain tissue microstructure is different from that of adult brain [36, 37]. Additionally, the fetal brain undergoes rapid development within a short period. Tissue microstructure undergoes significant changes during gestation as white matter tracts emerge, develop, and myelinate at varying time points and with different rates throughout this period [30, 34]. Thus, analyzing the fetal brain dMRI data requires dedicated computational methods that are not widely available. As a result, little is known about the development of the structural connectome in the fetal stage.

This represents a significant gap in knowledge as the fetal period is a dynamic and critical stage in brain development [3, 26, 38, 39, 40, 41]. Neurogenesis, neural migration, synapse formation, and axonal growth all begin before birth, forming the brain’s microstructure and laying the foundations of its networks [42, 43, 44, 45, 46, 3, 47, 27, 48, 49, 50]. It is well known that adult-like brain structures and a highly organized connectome develop in utero [46, 51, 49, 52, 34, 53, 54]. There is also growing evidence that the structural connectivity of the fetal brain can be disrupted by diseases and environmental factors, potentially resulting in lifelong neurodevelopmental and psychiatric disorders [39, 55, 56, 57]. For example, prenatal exposure to maternal stress, congenital heart disease, and brain malformations can alter the structural connectivity of the brain in utero [58, 59, 60, 61, 62, 63, 29, 64, 65, 66]. Therefore, a quantitative assessment of the structural connectome during the fetal period can deepen our understanding of the formation and development of the brain networks and may facilitate the identification of high-risk fetuses [8, 67, 28, 65, 68].

Prior studies on the fetal brain are limited in number and suffer from several limitations. First, they have analyzed relatively small populations [47, 65, 29] and have been restricted in terms of gestational age range considered [47, 65]. Moreover, they have often relied on basic and simplistic computational methods. For example, they have inferred the local orientation of major fibers using a diffusion tensor model, which cannot resolve crossing fibers [69, 65, 70]. This is an important limitation because patterns of fiber crossings are know to emerge early in the fetal stage [71]. Most prior works have used simple measures of connection strength, e.g., the number of streamlines, which are known to be inadequate and biased [11, 19, 12]. Additionally, prior works did not employ a true anatomically constrained tractography approach. This has been in part due to the fact that techniques for fetal brain segmentation and parcellation in dMRI have not been available. Such techniques based on deep learning have only recently become available [70, 69]. Hence, prior works usually did not include tissue segmentation, while for parcellation they relied on the registration of T2-weighted image atlases of fetal or neonatal brain [70, 69]. Unavailability of accurate tissue segmentation leads to very high rate of false positive streamlines because anatomical constraints cannot be enforced. On the other hand, registration of T2-weighted images to dMRI is highly challenging, with potentially high parcellation errors that significantly impact the connectivity analysis results. Furthermore, most prior studies have not used microstructure-informed filtering to improve the tractography results [72, 69]. Finally, because they rely on standard computational methods originally developed for adult brains, they are prone to suboptimal or erroneous tractography results [73, 74].

In this work we aimed to chart the development of structural connectivity in the fetal period. We used the recent release of the Developing Human Connectome Project (dHCP) dataset with dMRI scans from approximately 250 fetuses between 22 and 37 gestational weeks [75]. We leveraged fetal-specific methods for tissue segmentation, parcellation, and tractography. We computed the structural connectome for each fetus and determined how the connectivity metrics changed as a function of gestational age. We assessed the reproducibility of the results using bootstrapping methods. Moreover, we developed two methods to compute representative connectomes for each gestational week. One of these methods was based on precise spatial alignment and averaging of the imaging data, whereas the other method was based on aggregating the connectomes of individual subjects. We applied these methods to compute age-specific connectomes that depicted the temporal changes in structural connectivity between 22 and 37 gestational weeks. Using these atlases, we analyzed the major developmental patterns in intra- and inter-lobe connectivity within and across brain hemispheres. Overall, this study offers new insights into the development of the structural connectome in its earliest stages and provides benchmarks for the normative development of structural connectivity in the fetal period.

## 2 METHODS

### 2.1 Data and Preprocessing

This study used the fetal data from the dHCP dataset. The dataset includes 279 scans from 250 fetuses. We only considered scans with both structural (T2-weighted) and dMRI images, resulting in 259 MRI scan data from 239 fetuses. The dMRI data have been processed with the spherical harmonics and radial decomposition (SHARD) pipeline [75]. This method models the dMRI signal using a data-driven basis composed of spherical harmonics combined with radial decomposition, hence representing the dMRI signal across both angular and b-value dimensions without assuming a specific microstructural model. The framework also incorporates slice-level outlier detection, distortion correction, and slice profile modeling to down-weight or reject corrupted slices. The final output is a motion-corrected, reconstructed dMRI volume that can then be used for downstream analysis. We did not perform any further preprocessing on the data, except for registering the dMRI data to the T2-weighted image for each fetus and resampling both dMRI and T2-weighted images to an isotropic voxel size of 1 mm. Quality control was performed by an expert with six years of experience in fetal dMRI analysis based on careful visualization of (1) the diffusion tensor maps and fiber orientation maps in three orthogonal views, and (2) whole-brain tractograms in 3D. Of the 259 scans, 20 were excluded due to poor diffusion tensor or fiber orientation maps and 41 were excluded because of poor tractography. As a result, a total of 198 scans, each from a different fetus, were selected for structural connectivity analysis in this work. Figure 1 shows the histogram of gestational weeks for these fetuses. A total of 105 fetuses were male and 93 were female.

**FIGURE 1.**
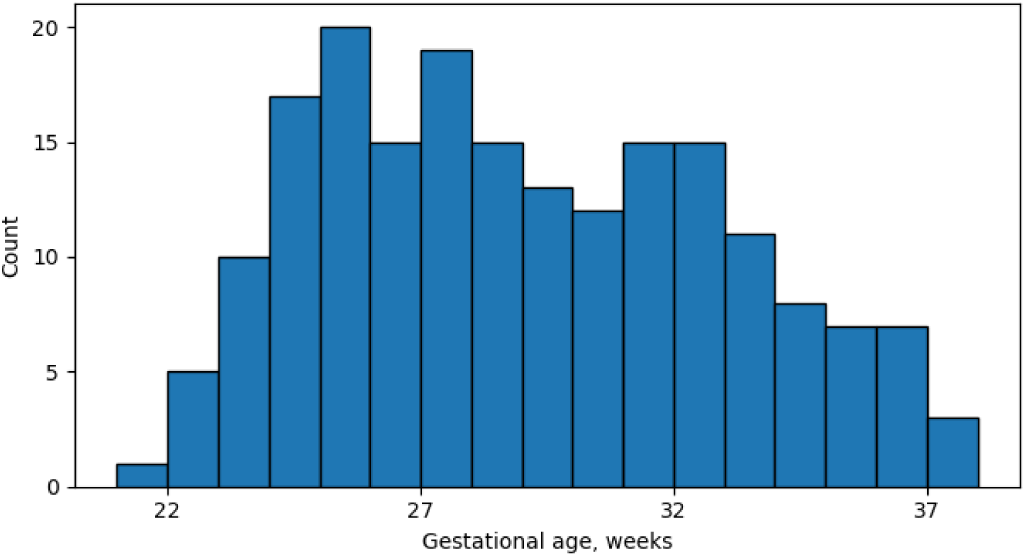
Histogram of the gestational ages of the 198 fetal subjects included in this study.

### 2.2 Fiber Orientation Estimation and Tractography

Each dMRI scan in this dataset consists of measurements with b=0 (n=15), b=400 (n=46), and b=1000 (n=80) *s* /*mm*^2^. As also reported in prior works [36], we found that standard multi-shell multi-tissue constrained spherical deconvolution (CSD) [76] was unable to correctly estimate the partial volume fractions for white matter, gray matter, and cerebrospinal fluid (CSF) in the fetal brain. Therefore, to estimate the fiber orientation density (FOD), we used the single-shell CSD [77], applied to the b=1000 shell data. We also computed the diffusion tensor images using the b=1000 *s* /*mm*^2^ data, which consistently produced visually superior results than with the b=400 *s* /*mm*^2^ shell data.

We applied a deep learning method [78], using the diffusion tensor image as input, to compute the tissue segmentation. Segmentation labels included white matter, cortical gray matter, subcortical gray matter, and CSF. For the cortical plate, we found that the segmentation from the T2-weighted image resulted in more accurate delineation of cortical foldings. Therefore, we combined the segmentation of the cortical plate from the T2-weighted image with segmentation of the white matter, sub-cortical gray matter, and CSF from dMRI. The same deep learning method [78] also parcellated the brain cortex into 88 regions, 44 in each hemisphere. Specifically, given the diffusion tensor image as input, the deep learning model computes a complete brain parcellation map. The definition of these regions are based on a publicly-available fetal brain atlas [79].

We performed anatomically constrained tractography using the iFOD2 algorithm [80] with a step size of 0.5 mm to propagate the streamlines, with a well-tested pipeline optimized for tractography of the fetal brain in the second and third trimesters [73]. Streamline seeding was performed on all white matter voxels and also on the boundary between white matter and gray matter. We observed that some of the tracts were more fully reconstructed by using the diffusion tensor as input while others were better reconstructed using the FOD computed with CSD. Moreover, a single setting of the angle threshold and FOD cutoff threshold was not adequate for computing all tracts. Therefore, we adopted an ensemble tractography approach by choosing ten different combinations of fiber orientation inputs and the angle / FOD cutoff thresholds and computing a separate initial tractogram with each of the ten combinations. Five of these tractograms were computed using the diffusion tensor as input and the remaining five were computed using the CSD-computed FOD. The angle threshold was varied in the range [10^◦^, 35^◦^], and FOD cutoff threshold was varied in the range [10^−5^, 10^−3^]. Table 1 provides detailed settings for this approach. Each of the initial tractograms included 10^5^ streamlines. We simply merged these tractograms to obtain the final tractogram with 10^6^streamlines. Note that the iFOD2 algorithm requires an FOD map as input. In order to apply this method with the diffusion tensor map, we computed its equivalent orientation distribution function using the following equation ODF(*u*) = 1/(4*π D* ^1/2^ (*u^T^ Du*)^3/2^), where *D* is the diffusion tensor and *u* is the arbitrary orientation [81, 82].

**TABLE 1.** Detailed settings used in the ensemble tractography approach. Abbreviations: FOD (fiber orientation density), DTI (diffusion tensor imaging), CSD (Constrained Spherical Deconvolution).

| Fiber orientation input | DTI | DTI | DTI | DTI | DTI | CSD | CSD | CSD | CSD | CSD |
| --- | --- | --- | --- | --- | --- | --- | --- | --- | --- | --- |
| FOD cut-off | $10^{-5}$ | $10^{-5}$ | $10^{-4}$ | $10^{-4}$ | $10^{-4}$ | $10^{-4}$ | $10^{-4}$ | $10^{-3}$ | $10^{-3}$ | $10^{-3}$ |
| Angle threshold (degree) | 10 | 20 | 25 | 30 | 35 | 15 | 20 | 25 | 30 | 35 |

For each brain, we computed the structural connectome using the approach described in [10, 83]. This approach uses the SIFT2 algorithm [84] to properly weigh the contribution of the streamlines to the structural connectivity strength. Note that SIFT2, as well as all subsequent computations, were applied on the complete tractogram that is obtained by merging the results of the 10 individual tractography runs. The connectomes were adjusted using the *µ* coefficient [10, 83].

### 2.3 Connectivity Metrics and Statistical Analysis

From the structural connectome of each fetus, we computed standard graph-theoretic descriptors of structural connectivity including global efficiency (GE), local efficiency (LE), characteristic path length (CPL), clustering coefficient (CC), and small-world index (SWI). All graph-theoretic metrics were computed on weighted structural connectomes using definitions presented in [13]. The weighted clustering coefficient for each node was computed using the formula by Onnela et al. [85]. The weighted characteristic path length was computed as the arithmetic mean of the shortest path distances, where the distance between two nodes was defined as the inverse of the edge weight, ensuring that stronger connections correspond to shorter path lengths. The small-world index was calculated as *σ* = (*C* /*C*_rand_)/(*L*/*L*_rand_) [86], where *C*_rand_ and *L*_rand_ are the mean clustering coefficient and characteristic path length of 100 random networks with the same degree distribution [87, 13]. We computed the strength of each node as the sum of connection strengths to all its neighbors. We performed linear regression analyses to determine whether the connectivity metrics, the connection strength of each node, and the strength of each connectome edge (i.e., connection between each node pair) changed significantly with gestational age.

### 2.4 Computation of Age-specific Connectomes

There is usually significant inter-subject variability in neuroimaging data. Atlases are used to characterize the population averages and represent typical brain development. We aimed to compute a representative structural connectome for each gestational week between 22 and 37, at one-week intervals. Specifically, we would like to leverage the data from individual fetuses around each gestational week to compute a connectome that represents the typical connectome for that gestational week. We refer to these as “age-specific connectomes”. We developed two different approaches for this purpose, one approach based on aggregating the connectomes computed for each individual fetus and another approach based on data averaging in the image space. These two approaches are described in detail below.

#### 2.4.1 Approach 1: Connectome aggregation

In this approach, we reconstructed an age-specific connectome for each gestational week between 22 and 37 based on the connectomes computed separately for individual fetuses. Each connectome (i.e., weighted adjacency matrix) is represented as a square symmetric matrix in **R***^d^* ^×^*^d^*, where *d* = 88 is the number of connectome nodes. Let us denote the set of connectomes for all *N* individual fetuses with 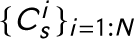, where 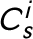 is the *subject* connectome for fetus *i*. Our goal is to compute a set of representative *template* connectomes 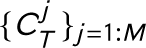, one for each of *M* gestational weeks. We consider three desiderata for the connectome template.

##### Desideratum 1-Representation

We would like the representative connectome for each week to closely represent the connectome for individual fetuses with a similar gestational age. To quantify the difference between the template and individual connectomes, we use the graph edit distance (GED). The GED between two graphs *C* ^1^ and *C* ^2^ is computed as the (minimum) cost of a set of edit operations to transform one graph into another [88]. These edit operations include insertion, deletion, and substitution of either a node or an edge. In general, for arbitrary graphs, computing the exact GED is Non-deterministic Polynomial-time hard (NP-hard). However, in this application all connectomes have the same fixed nodes. Therefore, GED only includes the costs associated with insertion, deletion, or substitution of edges. If we quantify the cost of each edge edit operation (i.e., edge insertion, deletion, or substitution) as the difference in edge weights due to that operation, GED will be the *ℓ*_1_ distance between the connectomes GED(*C* ^1^, *C* ^2^) = |*C* ^1^ − *C* ^2^ |. Therefore, we define the “representation loss” as 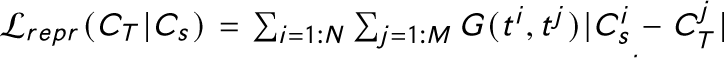. In this equation, *t ^i^*is the gestational age of subject *i* and *t*^j^ is that of the representative connectome 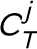. Moreover, *G* (*t ^i^*, *t ^j^*) = exp (− (*t ^i^* − *t ^j^*)^2^/2*σ*^2^) is a Gaussian kernel that encourages similarity of age-specific representative connectomes to the individual fetuses that are closest in gestational age. Similar to prior fetal imaging studies [79, 34], we set *σ* = 1.

##### Desideratum 2-Temporal consistency

We would also like to encourage temporal consistency in the computed age-specific connectomes. To this end, we penalize the GED between the templates with close gestational weeks. Similar to the representation loss above, we use a Gaussian kernel. Hence, our proposed loss term to encourage temporal consistency of the age-specific connectomes is 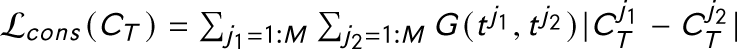.

##### Desideratum 3-Preservation of long connections

Another consideration is proportional representation/preservation of long-range connections. It is known that short-range connections are more likely to be shared across different structural connectomes [89, 90]. As a result, standard methods for combining connectomes based on connection consensus are biased towards preserving more of the short-range connections and discarding the long-range connections [89]. To compensate for this inherent bias, longer connections should be preferentially preserved in order to ensure that the template/average connectome is similar to the individual subject connectomes in terms of the distribution of connection lengths. One possible approach is to use a variable consensus threshold to preferentially preserve longer connections [89]. In this work, we use the Jensen-Shannon (JS) Divergence to penalize the difference between distance-wise distribution of the connectome weights. Specifically, for an arbitrary pair of connectomes *C* ^1^ and *C* ^2^, we compute the loss function L*_J S_* (*C* ^1^, *C* ^2^) = *D_J S_* (*p* (*C* ^1^), *p* (*C* ^2^)). Here, *p* (*C^i^*) denotes the distance-wise distribution of the connection weights for connectome *C^i^*, and *D_J S_* is computed based on the Kullback Leibler Divergence as *D_J S_* (*p*, *q*) = *D_K L_* (*p* ∥ (*p* + *q*)/2) + *D_K L_* (*q* ∥ (*p* + *q*)/2). We approximate *p* (.) using histograms with 20 bins. Moreover, we include a Gaussian weighting based on a similar logic as presented above for the other two loss terms. Hence, our proposed loss term is 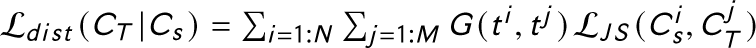.

Based on these three desiderata, we propose to compute the connectome templates by solving the following optimization problem:

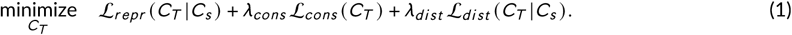

Note that in order to simplify the presentation, as in our descriptions above, in Equation (1) we have not shown the indices. To be clear, in Equation (1), 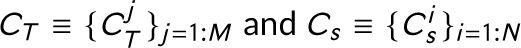. We solve a single optimization problem summarized in Equation (1) to recover age-specific connectomes for all *M* gestational weeks.

As the initial estimate to start the optimization in Equation (1), we computed *C_T_* via simple averaging of the connectomes in each gestational week. For example, for gestational age of 30 weeks we averaged the connectomes for fetuses between 29.5 and 30.5 weeks. We then iteratively optimized *C_T_* using stochastic gradient descent (SGD). In each optimization iteration we chose one of the subject connectomes 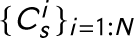 at random and optimized *C_T_* using SGD. We found that a small learning rate of 10^−5^ was necessary to ensure stable convergence of the optimization. Nonetheless, given the small size of the problem, the optimization was completed within a few seconds.

#### 2.4.2 Approach 2: Data averaging in the image space

The motivation for this approach is identical to the motivation for the connectome aggregation method above. Specifically, the goal is to average/aggregate the data from individual subjects to arrive at template connectomes that represent typical fetal brain connectomes at each gestational age. Technically, this approach is different as it performs the averaging in the image space. This approach is based on the precise alignment and “averaging” of the imaging data from subjects with a similar gestational age. Different steps of this approach are summarized in Figure 2. The main steps in this approach are summarized below.

**1.** Preprocess the structural MRI and dMRI data for each subject separately and register the dMRI to the structural MRI.
**2.** Divide all subjects into gestational age groups. In this work, we assigned each subject only to its closest gestational week. Specifically, all fetuses with a gestational age in the range [*t* − 0.5, *t* + 0.5) were assigned to the gestational week of *t*. For each gestational age group:

**a.** Compute the tissue response function for each subject [95] and average them to obtain a mean response function for the age group. Use this average response function to compute the FOD map for each subject using spherical deconvolution [77]. Compute a joint structural MRI and dMRI atlas for the age group via registration of data from all subjects within the age group. This is performed using both T2-weighted images and FOD maps to align the subject imaging data [92, 96]. The output of this step includes T2-weighted MRI, diffusion tensor, and FOD atlases for the gestational age group.
**b.** Compute the tissue segmentation and cortex parcellation with a deep learning method [78] using the diffusion tensor atlas as input. Use the cortical plate segmentation obtained from the T2-weighted atlas to refine the tissue segmentation and parcellation maps.
**c.** Compute the whole-brain tractogram based on the FOD and diffusion tensor atlases and tissue segmentation using our method described in [73]. We used an ensemble tractography approach similar to that described for individual fetuses in Section 2.2. Finally, compute the connectome.

**FIGURE 2.**
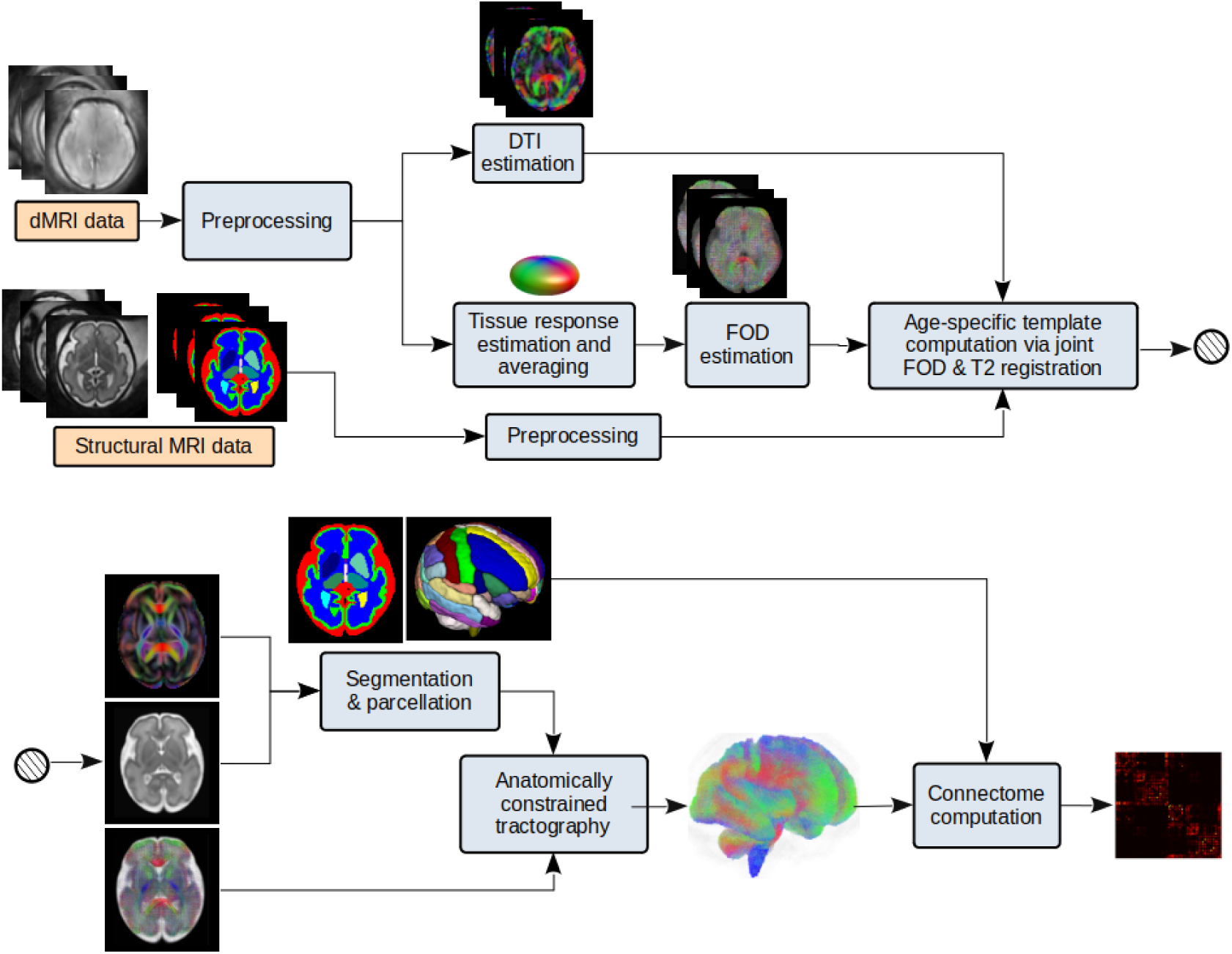
Image-space data averaging for computing a gestational age-specific connectome for each gestational week. Structural MRI and dMRI data from fetuses within a narrow window around the target gestational age are used as input to the pipeline. DTI and FOD maps are computed and precisely aligned using registration methods [91, 92, 93]. Segmentation/parcellation maps are computed either by fusion of subject-level data [94] or with deep learning methods [78].

We compared the above approaches with two existing techniques that are based on connectome aggregation [89, 90]: (1) Consensus-based thresholding, where the threshold is set such that the binary density of the connectome template is equal to the average binary density of the subject connectomes within the gestational age group. (2) Distance-preserved averaging, where different thresholds are used for different connection lengths such that the distribution of the connection lengths in the connectome template is close to that of the subjects. Detailed descriptions of these methods are presented in [89].

We assessed the different methods by comparing the age-specific representative connectomes to the connectomes of individual subjects in two ways:

**1.** First, we compared scalar connectivity metrics including GE, LE, CPL, and network strength between the representative connectomes and individual subject connectomes of the same age group.
**2.** Secondly, we evaluated the similarity of each individual fetal subject connectome to the connectome template of the corresponding age using a more sophisticated classification method similar to that proposed in [97]. In this approach each connectome was represented as a feature vector comprising all edge weights and eight topological descriptors for each node, including weighted degree, closeness centrality, betweenness centrality, eigenvector centrality, local efficiency, clustering coefficient, weighted number of triangles around the node, and PageRank [97, 98]. Pairwise differences between individual subject connectomes and template connectomes were summarized as the *ℓ*_1_, *ℓ*_2_, and *ℓ*_∞_ norms of the differences in their feature vectors. The resulting vector of size three is regarded as the feature vector that is used by a binary support vector machine classifier. The classifier is trained to predict whether a fetus and a template belonged to the same gestational age. We trained this method using a ten-fold cross-validation approach. Each time, we chose roughly 90% of the fetuses in each gestational week as training data. These fetuses were used to build the connectome templates and to train the support vector machine classifier. The trained classifier was then applied on the remaining held-out fetuses.

### 2.5 Reproducibility analysis

We evaluated the reproducibility of the reconstructed structural connectomes by assessing their consistency across independent subsets of dMRI measurements. We divided the *b* = 1000 measurements for each fetus into two non-overlapping halves and reconstructed a separate connectome from each subset. Denoting the connectomes reconstructed for fetus *i* from the two subsets as 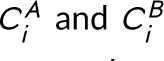, we computed the graph edit distance between them to quantify intra-subject variability. As a reference, we also computed inter-subject GEDs by comparing 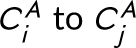 and 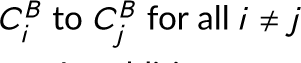.

In addition, we computed several graph-theoretical metrics – SWI, CPL, GE, LE, CC – from each connectome derived from the two measurement subsets. We then fitted linear models to assess how each metric varied with gestational age, separately for each subset. We assessed the differences in slope and intercept across the two data halves.

### 2.6 Network hubs

We identified the hubs of the structural connectome across gestational ages using three complementary centrality metrics: (1) degree centrality, which reflects the total connection strength of a node and captures its local importance; (2) betweenness centrality [99], which quantifies the extent to which a node lies on the shortest paths between other pairs of nodes, indicating its role as a connector or bridge; and (3) eigenvector centrality [100], which measures a node’s influence based on the importance of its neighbors and captures its global prominence within the network. While degree centrality is inherently a local measure, both betweenness and eigenvector centrality capture global aspects of the network.

## 3 RESULTS AND DISCUSSION

### 3.1 Development of the Structural Connectivity with Gestational Age

Figure 3 illustrates whole-brain tractograms and structural connectome matrices for four fetuses at 22, 27, 32, and 37 gestational weeks. All computational steps were visually inspected by an expert with six years of experience in fetal dMRI analysis. For comparison, Figure 4 presents age-specific average tractograms and connectomes, generated via data averaging in the image space as described in Section 2.4.2 and Figure 2.

**FIGURE 3.**
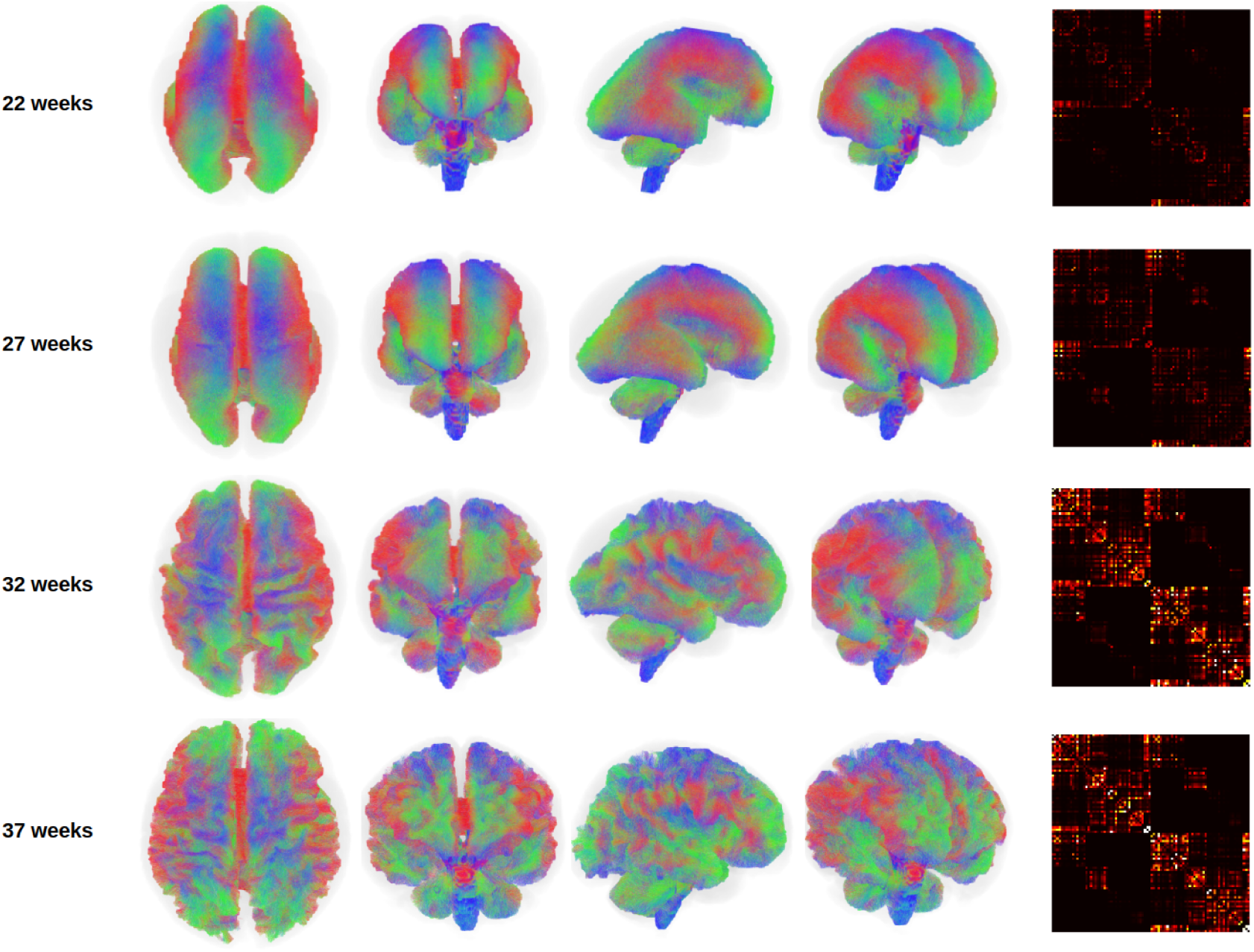
Different views of the whole brain tractograms, and the connectome matrices for four fetuses at different gestational ages. The tractograms for the lower gestational ages have been enlarged to the size of the higher gestational ages for better visualization. The connectomes for all gestational weeks have been shown with the same display intensity window of [0,5].

**FIGURE 4.**
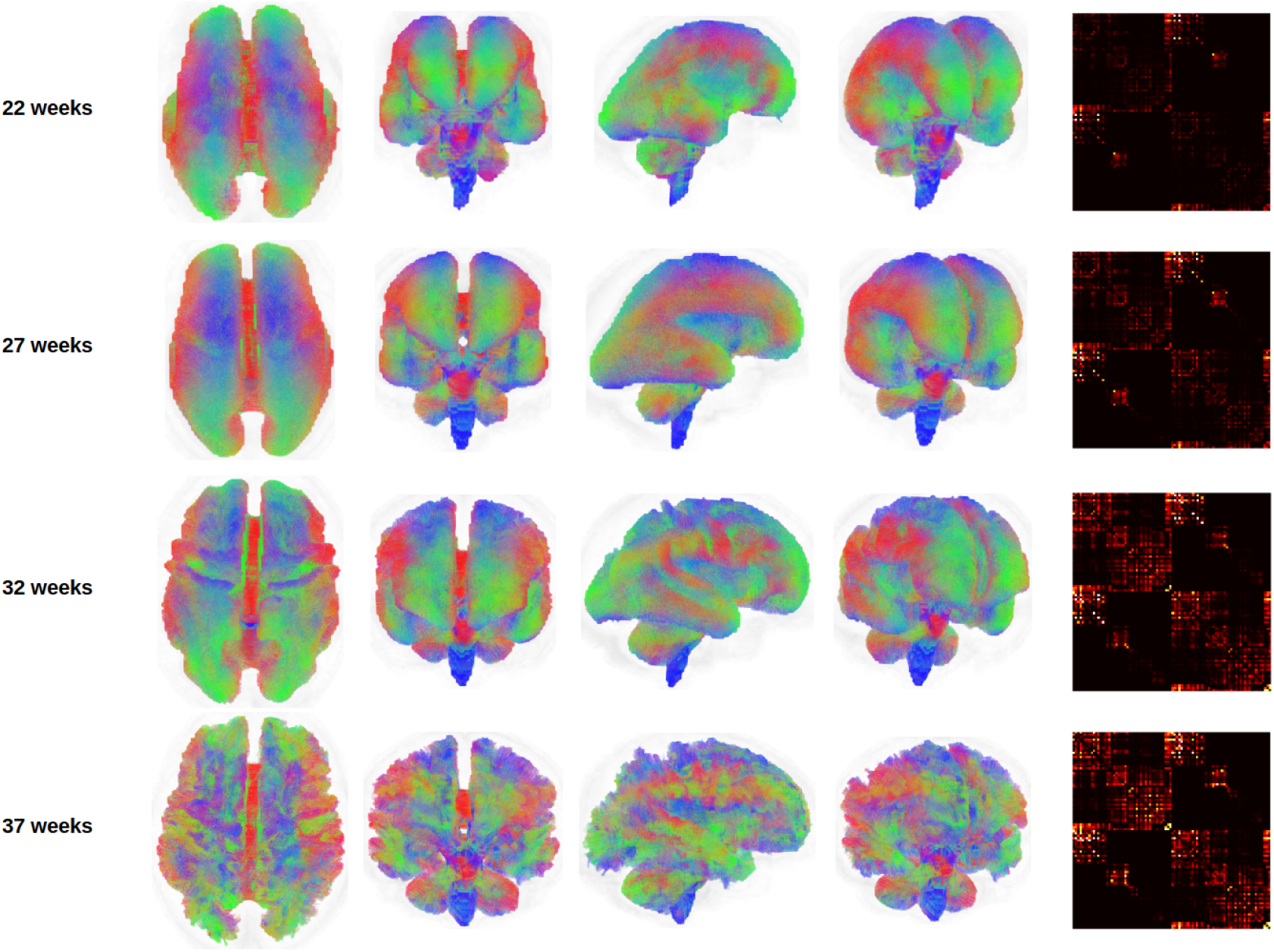
Tractograms and connectomes computed via data averaging in the image space, as described in Section 2.4.2, for four different gestational ages. The tractograms for the lower gestational ages have been enlarged to the size of the higher gestational ages for better visualization. The connectomes have been shown with the same display intensity window of [0,5].

#### 3.1.1 Changes in the global network metrics

Figure 5 shows the developmental trajectories of several key graph-theoretical metrics – GE, LE, CPL, CC, SWI, and total network strength (defined as the sum of nodal strengths across all regions) – for individual fetuses between 22 and 37 gestational weeks. In addition to individual trajectories, the figure includes these metrics computed from two connectome templates, constructed using the methods described in Section 2.4.

**FIGURE 5.**
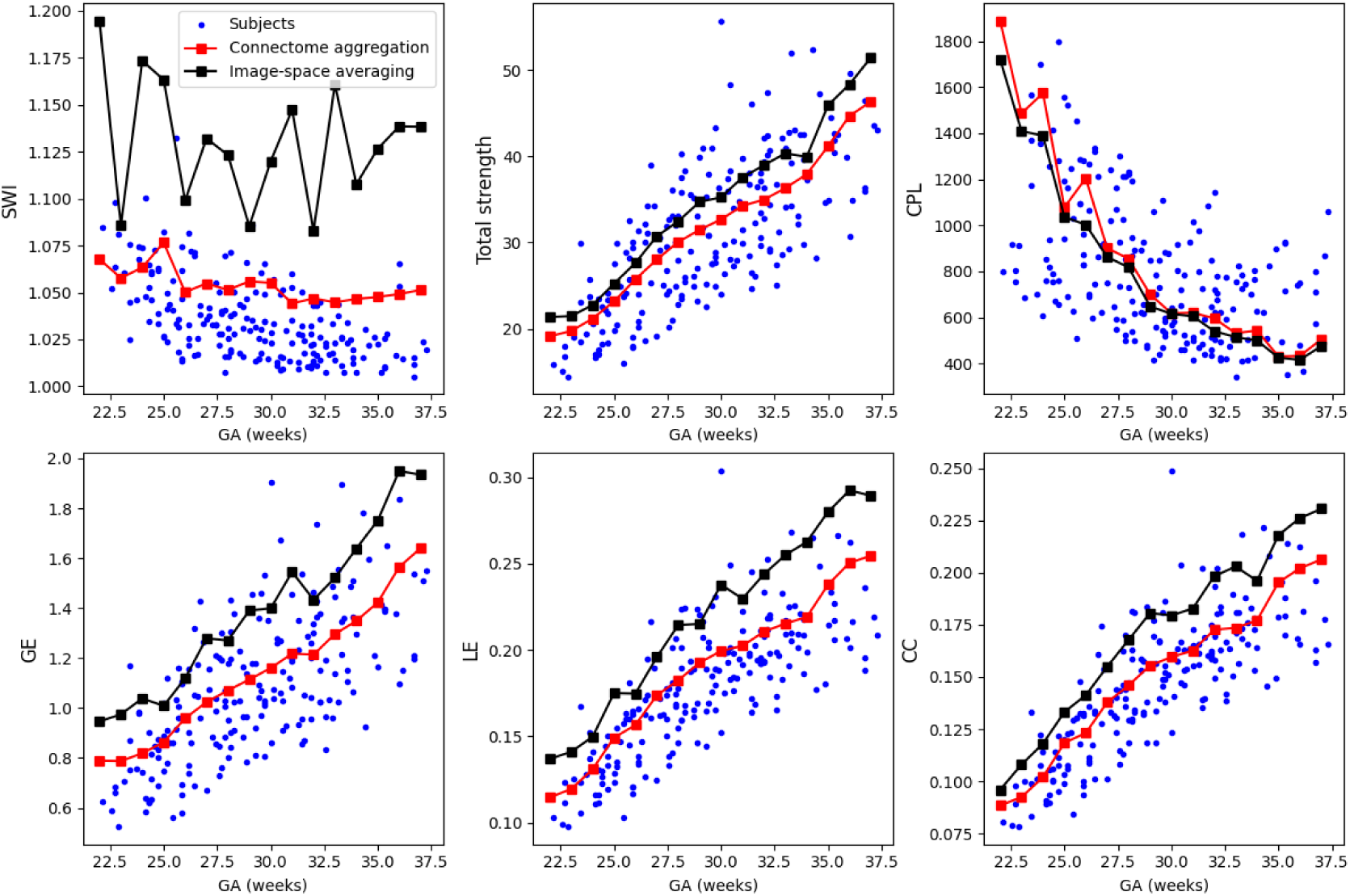
Plots of summary network metrics as a function of gestational age. Each plot shows the values for the subjects as blue dots and the connectome templates as red or black squares connected with line segments.

Overall, the plots reveal consistent and developmentally meaningful trends. Global efficiency steadily increases while characteristic path length decreases, indicating a progressive enhancement in network integration as gestation advances. These changes suggest that the fetal brain becomes increasingly optimized for long-range communication, enabling more efficient transfer of information between anatomically distant regions. Simultaneously, both local efficiency and the clustering coefficient increase, reflecting the gradual reinforcement of short-range, locally interconnected sub-networks that support specialized functional processing. The concurrent rise in integration (GE↑, CPL↓) and segregation (LE↑, CC↑) underscores the emergence of a small–world topology, an architectural hallmark of mature and efficient neural systems that balance local specialization with global integration.

A small-world configuration refers to a network architecture that supports both local specialization and global integration with high efficiency. It is defined by (1) a high clustering coefficient, reflecting densely interconnected neighboring brain regions that support local processing, and (2) a short characteristic path length, enabling rapid communication across distant brain areas [101]. In our analysis, SWI remains relatively stable across gestational ages, maintaining values in the range of 1.05–1.10. This stability occurs despite simultaneous increases in both global efficiency and clustering coefficient, suggesting that the relative balance between integration and segregation is preserved during late gestation. This observation implies that the fetal brain sustains a small–world topology throughout this period, supporting the hypothesis that small–world properties are intrinsic and robust features of neural systems emerging early in development and maintained as the brain undergoes structural maturation. The preserved SWI may also indicate that the underlying developmental mechanisms scale proportionally, affecting both local and global organization in a coordinated manner.

The SWI showed a relatively stable trajectory across gestation. SWI reflects the balance between normalized clustering and normalized characteristic path length rather than the absolute values of either metric. In our data, clustering increased while path length decreased with gestational age, and these coordinated changes appear to offset each other when compared with random networks, yielding a fairly constant SWI at the group level. Although the individual subject connectomes showed a slight downward trend in SWI at later gestational ages, this pattern was not fully captured by the connectome aggregation templates, which are designed to reflect the population-level balance of network organization. By contrast, the image-space averaging approach produced larger week-to-week fluctuations in SWI, likely because spatial alignment and voxel-wise averaging can alter fiber orientation information and the relative preservation of short-and long-range connections, making SWI more susceptible to displaying topological fluctuations.

Total network strength also shows a robust and monotonic increase with gestational age, reflecting a continuous rise in the overall connectivity of the brain. In this study, connection strength was quantified using the concept of fiber bundle capacity [10], a biologically informed metric that provides a more accurate estimate of white matter integrity and connection density than traditional measures such as streamline count. The steady increase in total strength likely mirrors the progressive development of white matter tracts, as axonal growth, myelination, and axon packing density all intensify throughout late gestation. Collectively, these findings highlight a phase of rapid and coordinated reorganization of the fetal brain’s structural network, marked by increasing complexity, stronger integration, and the gradual emergence of modular substructures. This evolving architecture is well aligned with the brain’s preparation for functional specialization and the demands of extra-uterine life.

Interestingly, the developmental curves computed from the connectome aggregation-based atlas closely mirror the average trends observed in the individual fetuses. This suggests that this method preserves the statistical and topological features of the underlying population well. In contrast, the connectome metrics computed from the template constructed via image-space averaging show notable deviations in the absolute values compared to those of individual subjects. Although the general trends with gestational age remain qualitatively consistent, the level shifts suggest that the averaging process in the image space introduces systematic alterations in the underlying connectivity patterns. This likely reflects the fact that when diffusion data from multiple fetuses (of similar gestational age) are aligned and averaged in the image space, the resulting atlas represents a spatial and microstructural average that smooths out some of the individual variability in structural connectivity. While such an atlas may offer a robust and anatomically coherent reference for the average brain at a given developmental stage, capturing shared morphological and diffusion features, it inevitably alters subject–specific details that contribute to individual connectome topology. Hence, the connectome templates obtained with this method may underrepresent inter–individual variability and distort the strength and specificity of white matter pathways. In particular, directional coherence, peak fiber density, or sharp anisotropic features that impact quantitative connectivity results may be blunted. These changes can systematically influence network topology, leading to systematic biases in graph–theoretical metric values that diverge from those observed in individual data.

#### 3.1.2 Changes in the nodal and edge strength with gestational age

We plotted the nodal strength versus gestational age for each of the 88 brain regions included in the connectome and observed that the nodal strength increased significantly as a function of gestational age for all 88 nodes. Linear regression analysis confirmed a statistically significant positive slope for all nodes, even under stringent control for multiple comparisons. Specifically, we used a significance threshold of *α* = 0.01 and applied Bonferroni correction to account for testing across 88 regions. Figure 6 shows the nodal strength at the earliest and latest gestational weeks considered in this work as well as the slope of increase in the nodal strength between 22-37 gestational weeks.

**FIGURE 6.**
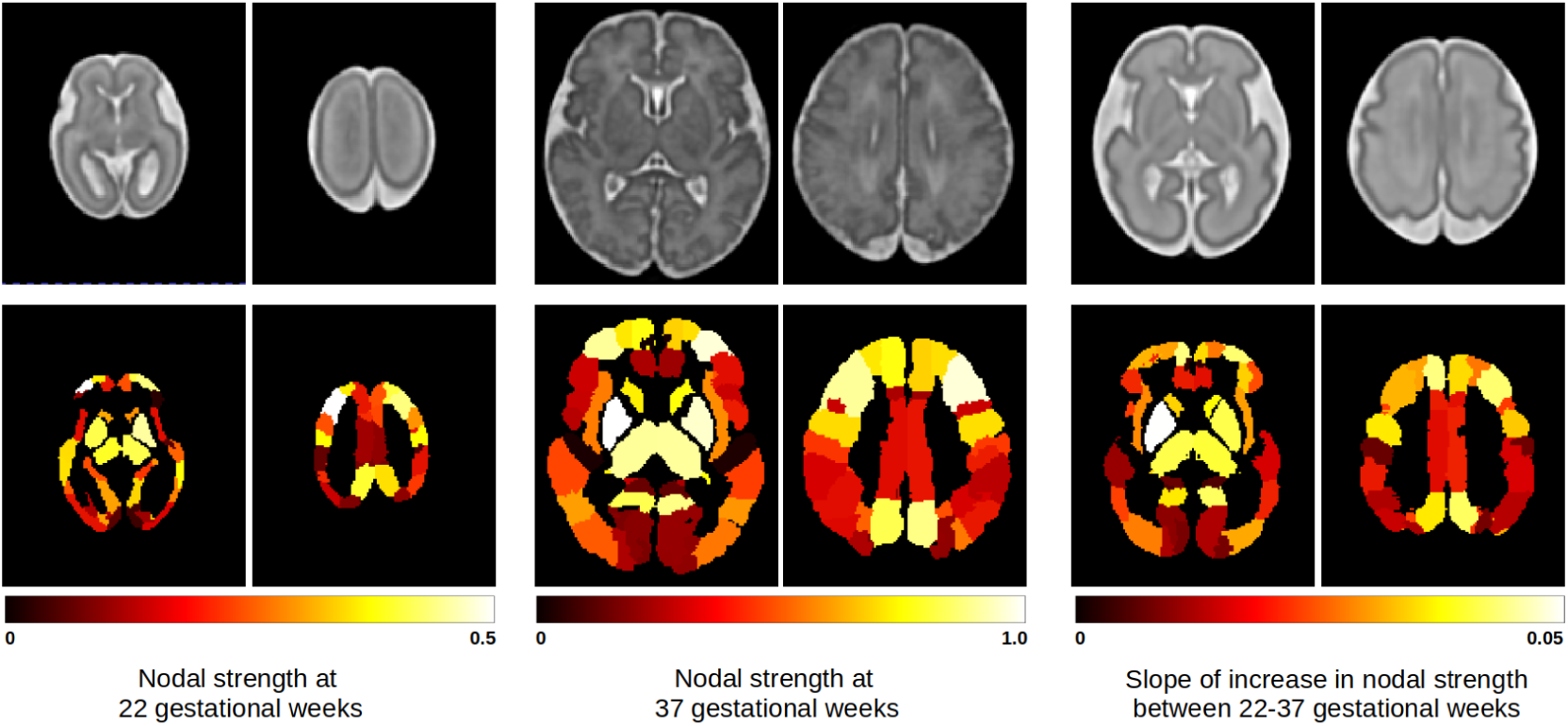
Axial slices of T2-weighted fetal brain atlases (top row) and the corresponding slices showing the nodal strength for different brain regions (bottom row). The two left-most columns show the nodal strength at the earliest age considered in this study (22 gestational weeks) and the two middle columns show the nodal strengths at the latest age (37 gestational weeks). The two right-most columns show the slope of the linear fit, displaying the rate of change, per week, in the strength for each node between 22-37 gestational weeks. The atlas template used in the right-most columns is for 30 gestational week.

We further investigated how the strength of individual connections between brain regions changed over gestation by analyzing the temporal changes in edge strength. To assess the reproducibility of these developmental trends, we employed a robust resampling strategy. Specifically, for each network edge, we randomly selected 75% of the subjects without replacement and tested for a statistically significant linear association between edge strength and gestational age. This procedure was repeated 100 times per edge, and for each repetition, we recorded whether the edge exhibited a significant change. As shown in Figure 7, we summarized the results by calculating the percentage of repetitions in which each edge showed a significant association. To ensure high robustness, we retained only those edges that were significant in at least 95% of the repetitions. Based on this criterion, 162 edges demonstrated a consistent increase in strength over gestation, while 12 showed a consistent decrease. The strengthening edges encompassed both short-range and long-range connections, linking neighboring regions within lobes as well as distant regions across lobes and hemispheres. Notably, robust increases were observed in connections involving the anterior, middle, and posterior cingulate gyri, as well as in interhemispheric connections between the hippocampi. In contrast, the few connections that weakened with gestational age were primarily interhemispheric, including those linking the postcentral gyri, superior parietal lobules, and inferior parietal lobules in the parietal lobe, as well as the superior and medial frontal gyri in the frontal lobe. These findings suggest that while most structural connections strengthen during late gestation, a small subset of connections may be selectively pruned or refined as part of the maturation process. Figure 8 shows the percentage change in the connection strength between 22 and 37 gestational weeks. Overall, the patterns observed in this figure are similar to Figure 7, with a larger number of strengthening inter-hemispheric connections showing up in Figure 8. Figure 8 also shows a widespread increase in connection strength with a much smaller number of weakening connections, where edges experiencing the greatest change in strength mostly overlap with those that found to change consistently in Figure 7.

**FIGURE 7.**
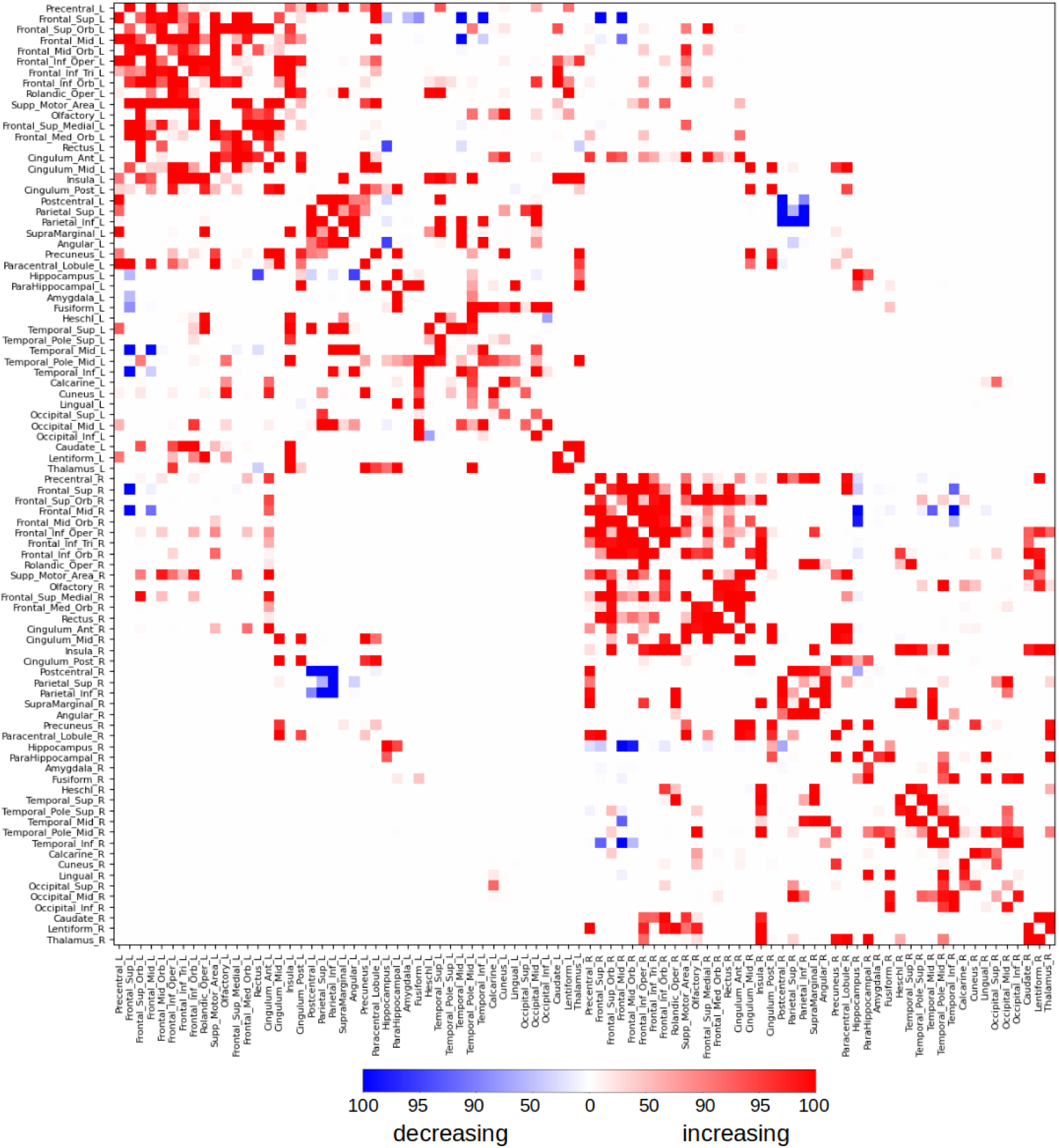
Change in the edge connection strength, i.e., the strength of connection between pairs of nodes, as a function of gestational age. We randomly selected 75% of the subjects and determined if there was a significant linear relation between the edge strength and gestational age. We repeated this 100 times for each edge. This figures shows the percentage of times the connection showed a significant increase (red) or decrease (blue) with gestational age.

**FIGURE 8.**
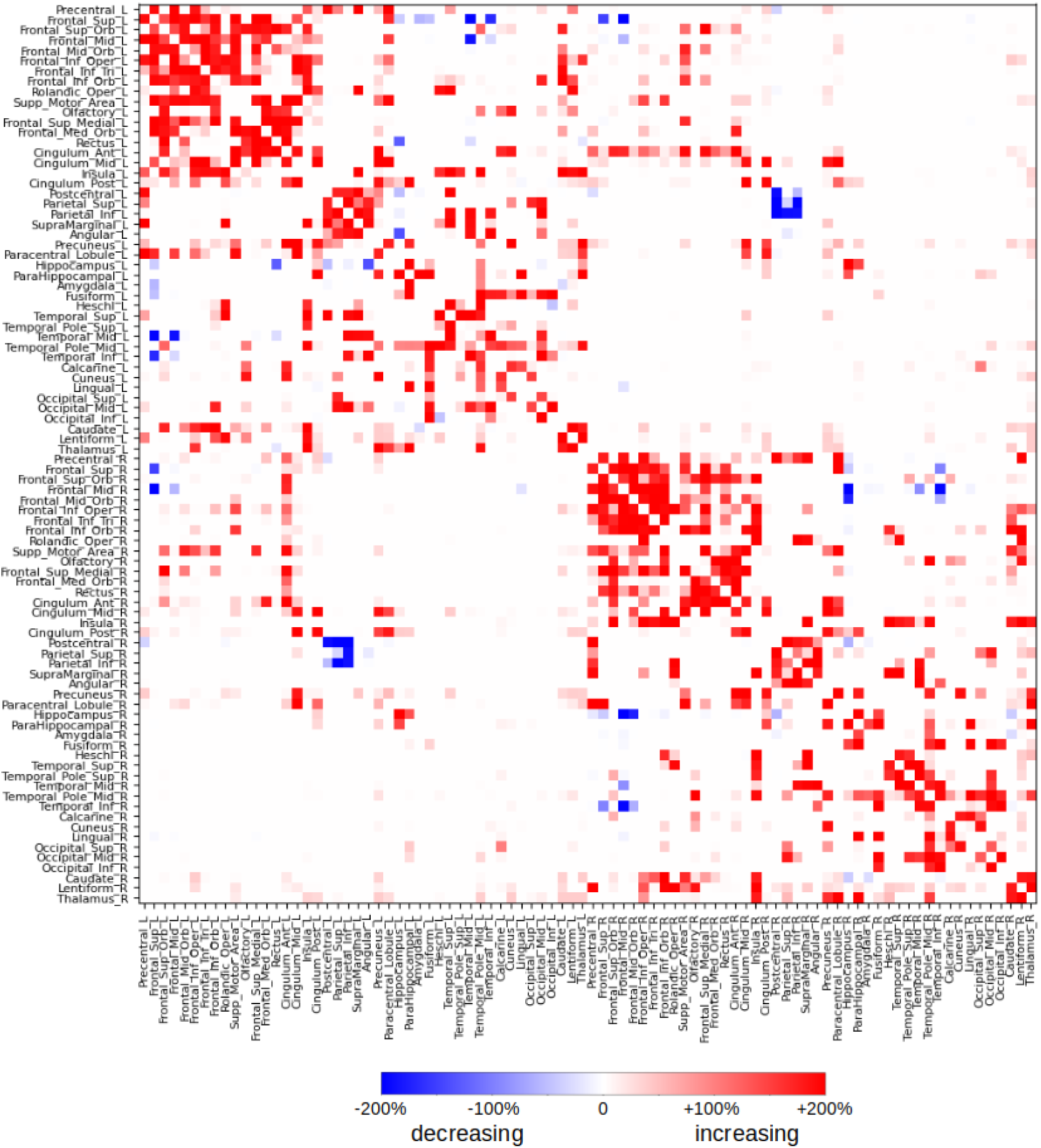
Percentage change in the strength of different connection edges between 22-37 gestational weeks.

To further investigate whether the magnitude of connection strengthening differed systematically between short-and long-range pathways, we analyzed the relationship between inter-nodal distance and the rate of connection strength change over gestation. As shown in Figure 9, there was a significant negative correlation between inter-nodal distance and the rate of connection strength change (p<0.001). Although most connections, both short-and long-range, showed positive strengthening over gestation, short-range connections exhibited significantly greater strengthening than their longer-range counterparts. This pattern indicates that the observed increases in global efficiency and decreases in characteristic path length (Figure 5) are not driven exclusively by long-range connections. Rather, the pronounced consolidation of short-range circuitry enhances local efficiency and clustering, while the concurrent, albeit more modest, strengthening of long-range connections supports global integration. Thus, local circuit refinement and global integration proceed in parallel during the fetal period. Moreover, examination of the distribution of the connection strength changes revealed a bimodal pattern: a two-component Gaussian mixture model provided a better fit than a single Gaussian (lower Bayesian information criterion), indicating that the edges naturally separate into two groups with distinct rates of strength increase. While connections in both groups showed strengthening over gestation, the cluster with larger changes was enriched with short-range edges and the cluster with more modest changes comprised primarily of long-range connections.

**FIGURE 9.**
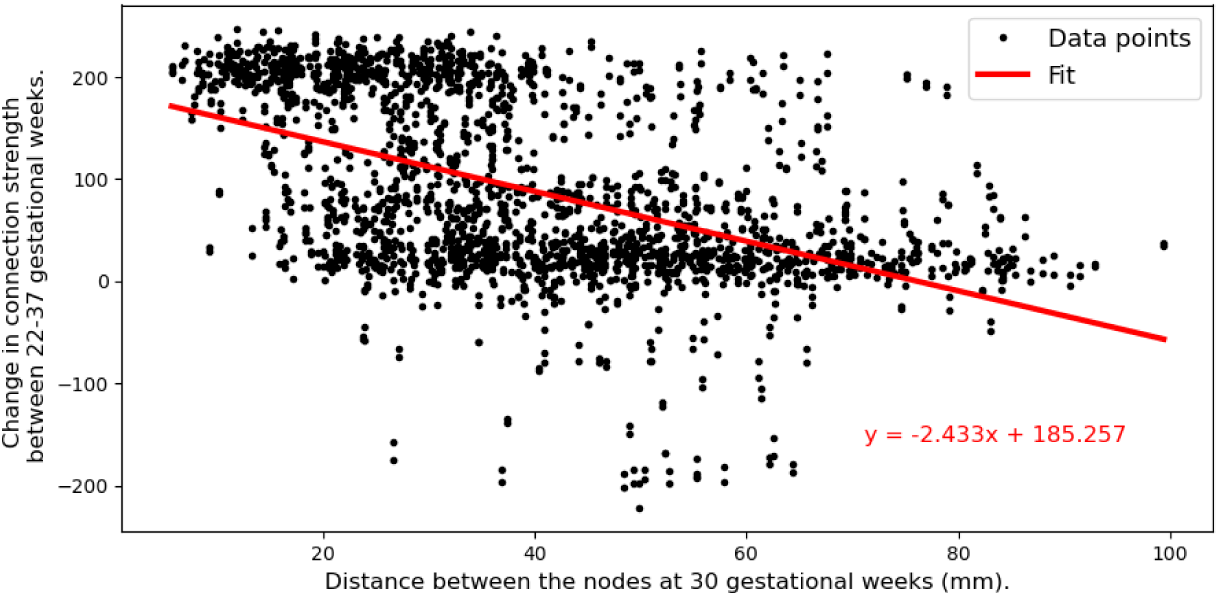
Relationship between inter-nodal distance and connection strength change over gestation. Each point represents a single edge (connection) in the connectome. The x-axis shows the Euclidean distance between the centroids of the two connected nodes, computed at 30 gestational weeks to account for brain growth. The y-axis shows the percentage change in connection strength between 22-37 weeks; positive values indicate strengthening over gestation. The solid line represents the linear fit, with a statistically significant negative slope (*p* < 0.001, *R* ^2^ = 0.258). Short-range connections (left side of the plot) show greater degree of strengthening, while long-range connections (right side) also strengthen but to a lesser extent.

#### 3.1.3 Network hubs across the gestation

Figure 10 illustrates the distribution of network hubs at four gestational ages, based on eigenvector centrality. Several brain regions consistently emerged as hubs across most gestational weeks (22–37 weeks), including the precentral gyrus, superior and middle frontal gyri, precuneus, lentiform, and thalamus. Some regions showed increasing prominence with gestational age, notably the supplementary motor area, medial portion of the superior frontal gyrus, and insula, increasing from approximately 0.10-0.12 to 0.15-0.17. Conversely, a few regions, such as the middle temporal gyrus and inferior temporal gyrus, exhibited a decline in hub status over time, decreasing from approximately 0.16-0.19 to 0.14-0.16, although they remained strongly connected.

**FIGURE 10.**
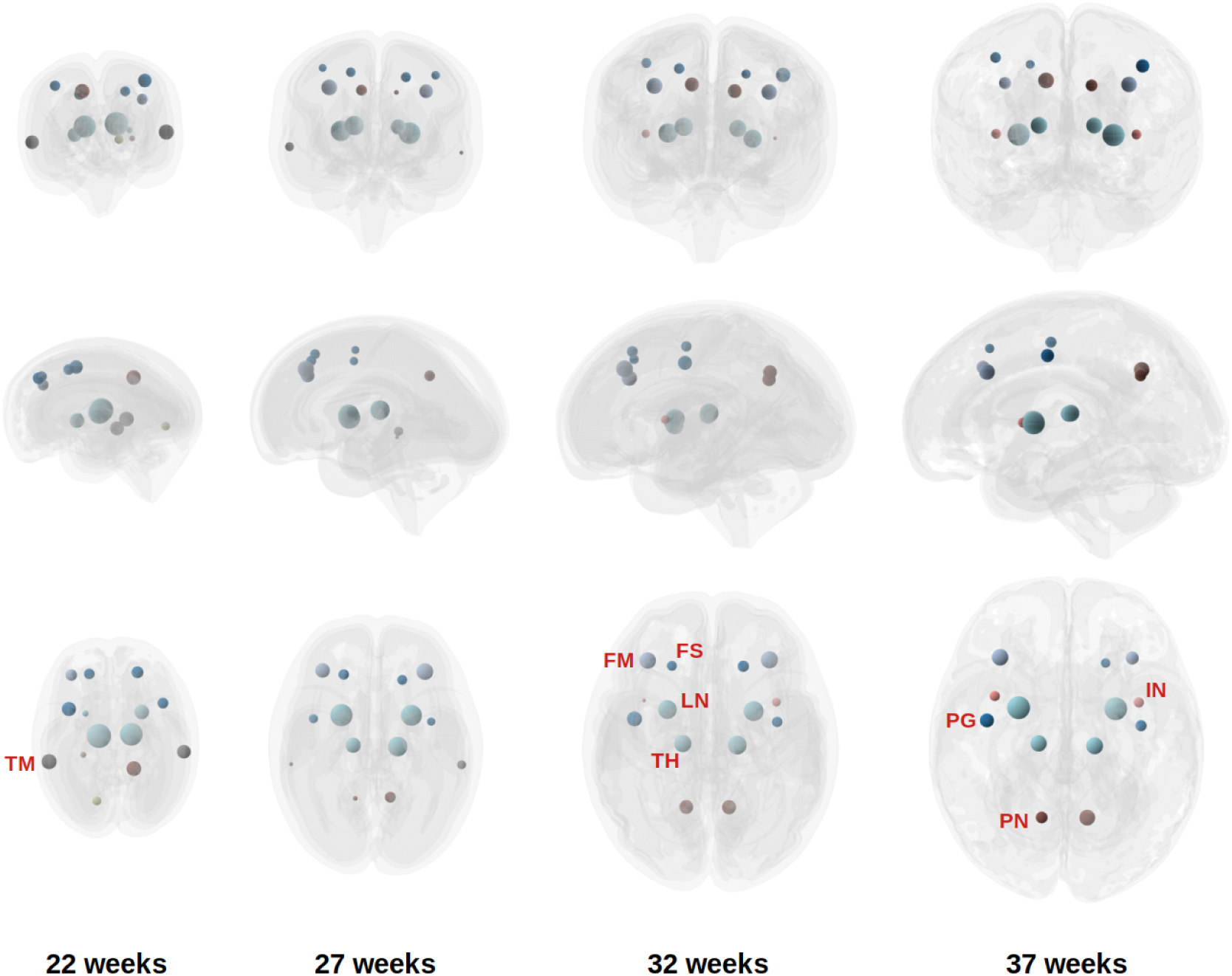
Network hubs identified on individual fetuses, averaged across all fetuses at each gestational week. The size of the sphere is proportional to the node strength for each hub. FS: Frontal superior gyrus; FM: Frontal middle gyrus; IN: Insula; LN: Lentiform; PN: Precuneus; PG: Precentral gyrus; TH: Thalamus; TM: Middle temporal gyrus.

Overall, there was substantial concordance among the three centrality measures, though some differences were observed. When defined by degree and betweenness centrality, the middle frontal gyrus and inferior temporal gyrus appeared less prominent, while the angular gyrus, fusiform gyrus, superior parietal lobule, and inferior parietal lobule emerged as more prominent hubs under these criteria.

Our findings broadly align with previous studies on structural connectome hubs in the human brain. In adults, commonly reported hub regions include the precuneus, superior frontal and superior parietal cortices, anterior and posterior cingulate cortex, insular cortex, temporal cortex, lentiform/putamen, thalamus, and hippocampus [102, 103, 104, 105, 106, 107, 108]. Several studies have also examined hub development in neonates and infants, often using functional MRI–based connectomes [109, 110, 48, 111]. Reported structural hubs in neonates include the dorsal and medial frontal cortex, parietal cortex, precuneus, hippocampus, insula, and anterior cingulate cortex [112, 8, 113].

Functional hubs during the fetal and neonatal periods have been reported to include the cingulate cortex, precentral and postcentral gyri, superior parietal lobule, cerebellum, angular gyrus, fusiform gyrus, insula, and association cortices near the fusiform facial area and Wernicke’s area [109, 114, 48, 111, 115, 116, 117, 46]. The considerable overlap between hubs identified in our study and those reported in the literature lends support to the biological plausibility and generalizability of our findings. Notably, however, we did not identify the hippocampus or the cingulate cortex as major hubs, despite their frequent appearance as hubs in prior studies. These discrepancies may reflect differences in imaging modality, gestational age range, or methodological approach, including the definition of hub metrics and tractography techniques. In fetal imaging, in particular, there is a lack of standard computational methods for cortical parcellation, and differences in the definitions and the extent of cortical regions can lead to variabilities in the determination of structural connectivity hubs.

Our findings also offer an opportunity to compare these dMRI-based results with the existing hypotheses regarding the emergence and maturation of brain network hubs, which have been proposed based on studies in model organisms, postmortem histology, and postnatal cohorts [118, 119, 112, 120]. A central tenet of this literature is that hub regions emerge prior to non-hubs and form a stable structural scaffold that guides subsequent network development [118]. Our data support this hypothesis. The spatial topography of structural hubs, as identified by eigenvector centrality and other metrics, showed consistency across the 22-37 week gestational window (Figure 10). Regions such as the precuneus, superior frontal gyrus, and thalamus were consistently identified as hubs from the earliest time points in our study, suggesting that the anatomical identity of core network elements is established by the early third trimester. This stability aligns with the proposition that the binary topology of the connectome is established prenatally, with subsequent development involving the strengthening and refinement of existing connections [118].

The mechanisms proposed to explain hub formation include the “old-gets-richer” principle, whereby early-born nodes accumulate connections from later-developing regions [119], and the concept of developmental heterochronicity along a sensorimotor-association axis [120]. Our data are consistent with both models. The progressive strengthening of hub connections that we observed throughout gestation (Figures 5 - 8) can be interpreted as the ongoing consolidation of an early-established structural core. Furthermore, the few connections that showed relative weakening over gestation were predominantly interhemispheric parietal and frontal connections, a pattern that may reflect early stages of the activity-dependent pruning and refinement processes that are known to sculpt cortical circuitry during development [42]. This selective weakening, alongside widespread strengthening, is compatible with the view that hubs form a stable “rich club” that serves as a high-capacity backbone, while peripheral connections undergo more dynamic reorganization [102, 112].

Our findings also align with prior work examining the developmental trajectory of rich-club organization in the preterm period. Ball et al. demonstrated that rich-club architecture is already present by 30 weeks gestation and that the principal development between 30 and 40 weeks involves a proliferation of “feeder” connections between core hubs and the rest of the cortex, rather than changes in the core connections themselves [112]. While our study spans a slightly different gestational window (22–37 weeks) and employed a different parcellation scheme, our results are broadly congruent. We observed robust increases in overall network strength and global efficiency, with pronounced changes occurring in connections involving hub regions. Although we did not perform a formal longitudinal analysis of feeder versus core connections, the pattern of widespread strengthening in hub-related edges is consistent with the view that the rich-club backbone is established early and subsequently becomes increasingly integrated with peripheral regions [112]. Together, these convergent findings from independent datasets and methodologies underscore the robustness of the developmental principles governing the emergence of human brain network architecture.

### 3.2 Age-specific Connectome Templates

#### 3.2.1 Comparison of different template reconstruction methods

Figure 11 presents the age-specific connectome templates for gestational weeks 22 to 37, generated using the proposed connectome aggregation method described in Section 2.4.1. For comparison, Figure 12 displays the corresponding templates constructed via image–space averaging, as detailed in Section 2.4.2. Both approaches yield connectomes that demonstrate a consistent increase in connection strength over gestation – within individual brain lobes, across different lobes, and between hemispheres. While the overall topological trends are broadly similar between the two methods, the templates derived from image–space averaging tend to show stronger inter-hemispheric connections. This is likely due to the spatial smoothing effect inherent in voxel-wise averaging across multiple subjects.

**FIGURE 11.**
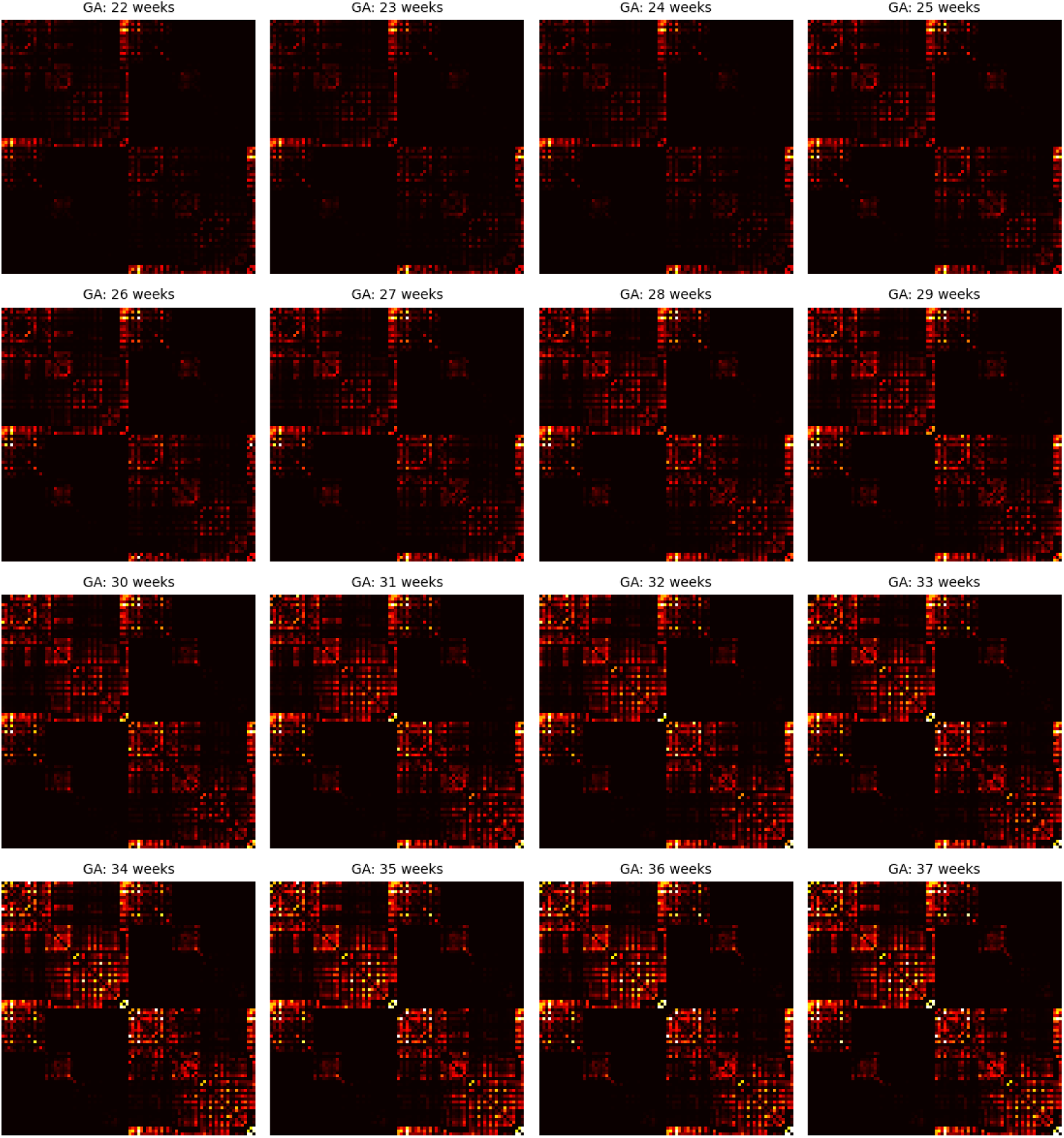
Connectome templates computed for every gestational age between 22 and 37 weeks with the connectome aggregation method proposed in Section 2.4.1.

**FIGURE 12.**
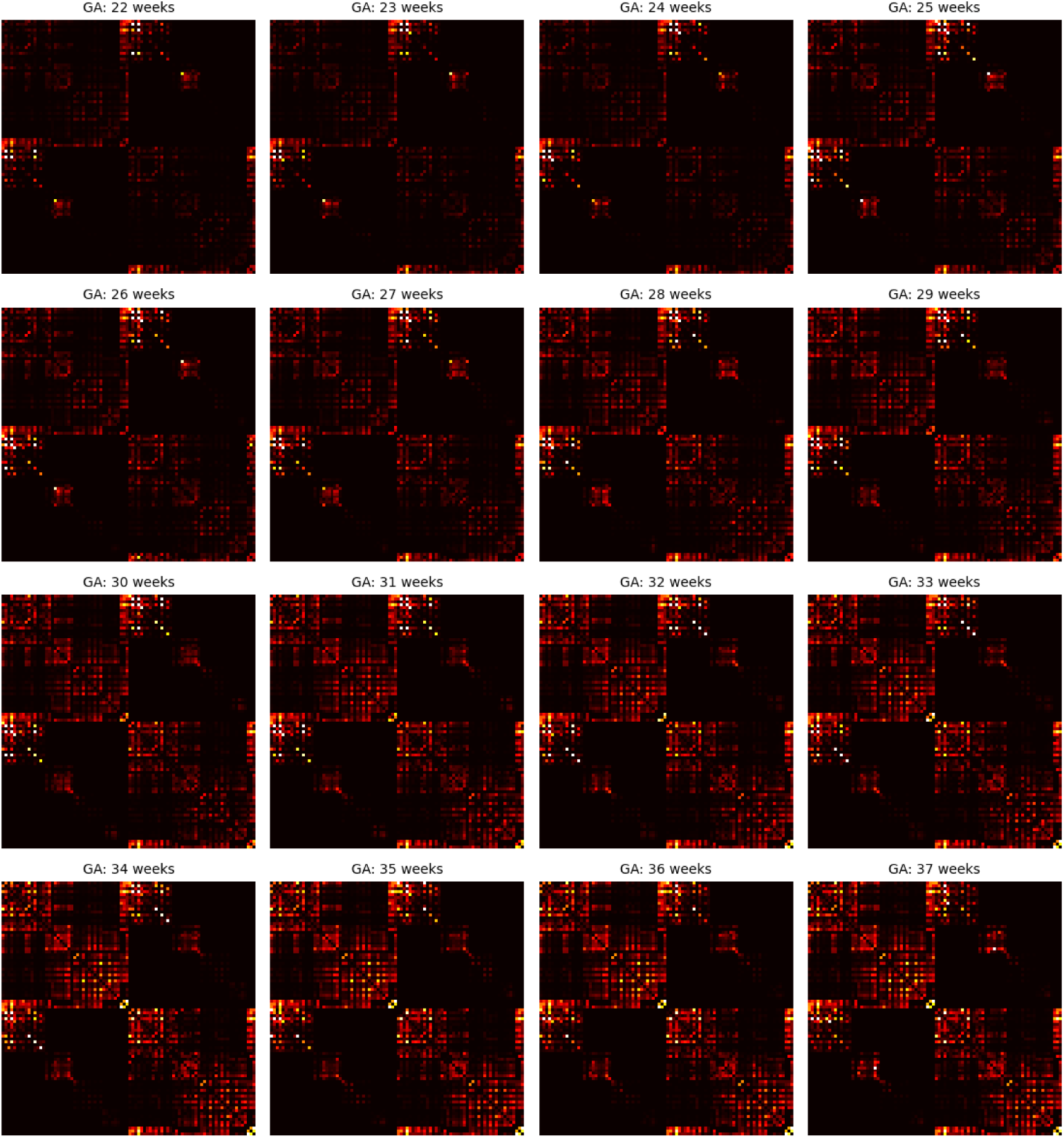
Connectome templates computed for every gestational week via data averaging in the image space described in Section 2.4.2.

To quantitatively evaluate the performance of these methods, we compared the resulting templates to the individual connectomes of age-matched fetuses. Table 2 summarizes this comparison, which also includes the baseline method of distance-preserved averaging of connectomes [89]. The results for consensus-based thresholding were largely similar to distance-preserved averaging and, hence, are omitted from this table.

**TABLE 2.** Quantitative comparison of the connectome templates computed using different methods. The first four rows show the difference in the connectivity metrics between the individual subjects and their age–matched connectome templates. The next two rows show the accuracy of predicting the age of an individual fetus based on the similarity of his/her connectome with the age–specific connectome templates. More specifically, these two rows show the error (in weeks) in the predicted age of individual fetuses based on the similarity to the connectome templates. The last row shows the overlap of the 10 top hubs between each fetus and its age–matched connectome. Abbreviations: GE (global efficiency), LE (local efficiency), CPL (characteristic path length), GED (graph edit distance).

|  | Connectome aggregation method | Data averaging in the image space | Distance-preserved averaging [89] |
| --- | --- | --- | --- |
| GE | $0.240 \pm 0.207$ | $0.375 \pm 0.422$ | $0.352 \pm 0.228$ |
| LE | $0.039 \pm 0.029$ | $0.058 \pm 0.036$ | $0.040 \pm 0.037$ |
| CPL | $246 \pm 228$ | $254 \pm 214$ | $256 \pm 126$ |
| Network strength | $6.63 \pm 6.21$ | $8.05 \pm 6.83$ | $8.38 \pm 6.40$ |
| Age prediction error (in weeks), computed using on a nearest-neighbor classifier based on graph edit distance (GED) | $0.98 \pm 0.95$ | $2.01 \pm 1.39$ | $2.77 \pm 1.45$ |
| Age prediction error (in weeks), computed using a support vector machine classifier as in [97] | $0.90 \pm 0.81$ | $1.44 \pm 1.37$ | $1.46 \pm 1.85$ |
| Hub overlap | $0.83 \pm 0.08$ | $0.80 \pm 0.06$ | $0.72 \pm 0.07$ |

For each template, we measured the difference in several key graph–theoretical metrics including GE, LE, CPL, and total network strength, between the template and the corresponding individual connectomes. Across all metrics, the connectome aggregation method yielded templates that more closely resembled individual subject connectomes. Figure 5 shows a visual representation of these metrics.

We also evaluated the ability of each connectome template to capture age-related patterns using classification experiments. First, a nearest-neighbor classifier based on GED was used to predict the gestational age. Templates constructed via connectome aggregation achieved the lowest prediction error (0.98 ± 0.95 weeks), substantially out-performing both image–space averaging and element–wise averaging, which yielded errors greater than 2 weeks. We should point out, however, that this evaluation may be biased towards the connectome aggregation method. This is because the GED was used in the optimization formulation for reconstructing these templates, as described in Section 2.4.1. Next, we implemented a more sophisticated classification framework based on the method in [97], described in Section 2.4. This method achieved an error of 0.90 weeks with the connectome aggregation templates, outperforming the other two methods (1.44 and 1.46 weeks, respectively). All results were validated using ten-fold cross-validation.

Finally, we assessed how well each method preserved the location of high-centrality nodes (i.e., hubs). For each gestational age, we identified the top 10 hubs in individual fetuses and their corresponding templates, and calculated the percentage of overlap. As shown in Table 2, the connectome aggregation–based templates had the highest agreement with the individual–level hubs (83% overlap), followed by image–space averaging (80%), and distance-preserved averaging (72%). These results indicate that connectome aggregation not only preserves overall network topology but also retains region–specific nodal prominence better than the other methods.

A general consideration when constructing any template or atlas is that the resulting representation inherently smooths over inter-individual variability to capture central tendencies of the population. Our connectome aggregation method is no exception; the optimization framework encourages temporal consistency and preservation of long-range connections, which may downweight genuine but idiosyncratic features present in individual fetuses. This trade-off is inherent to any template-building approach, whether based on connectome aggregation or image-space averaging, and reflects the fundamental purpose of an atlas: to represent the population norm rather than any single subject. We have attempted to mitigate this bias by designing loss functions that prioritize fidelity to individual data (i.e., the representation term in our optimization formulation) and by validating our templates against multiple criteria, including similarity to individual connectomes and age prediction accuracy. Nevertheless, these templates are optimized to represent population trends, and template-based analyses should be complemented with subject-level investigations to fully capture the range of individual variations.

#### 3.2.2 Analysis of spatiotemporal trends based on the connectome templates

As summarized in Table 2 and illustrated in Figure 5, the connectome templates reconstructed using the connectome aggregation method outperformed those generated by alternative approaches in preserving key network characteristics. These high–fidelity templates were therefore used to analyze the developmental evolution of structural connectivity. Figure 13 presents four sets of plots illustrating the temporal dynamics of connectivity across different brain regions computed using these templates. Consistent with these visualizations and the connectome matrices shown in Figure 11, the fetal brain undergoes marked and progressive changes in structural connectivity between 22 and 37 gestational weeks.

**FIGURE 13.**
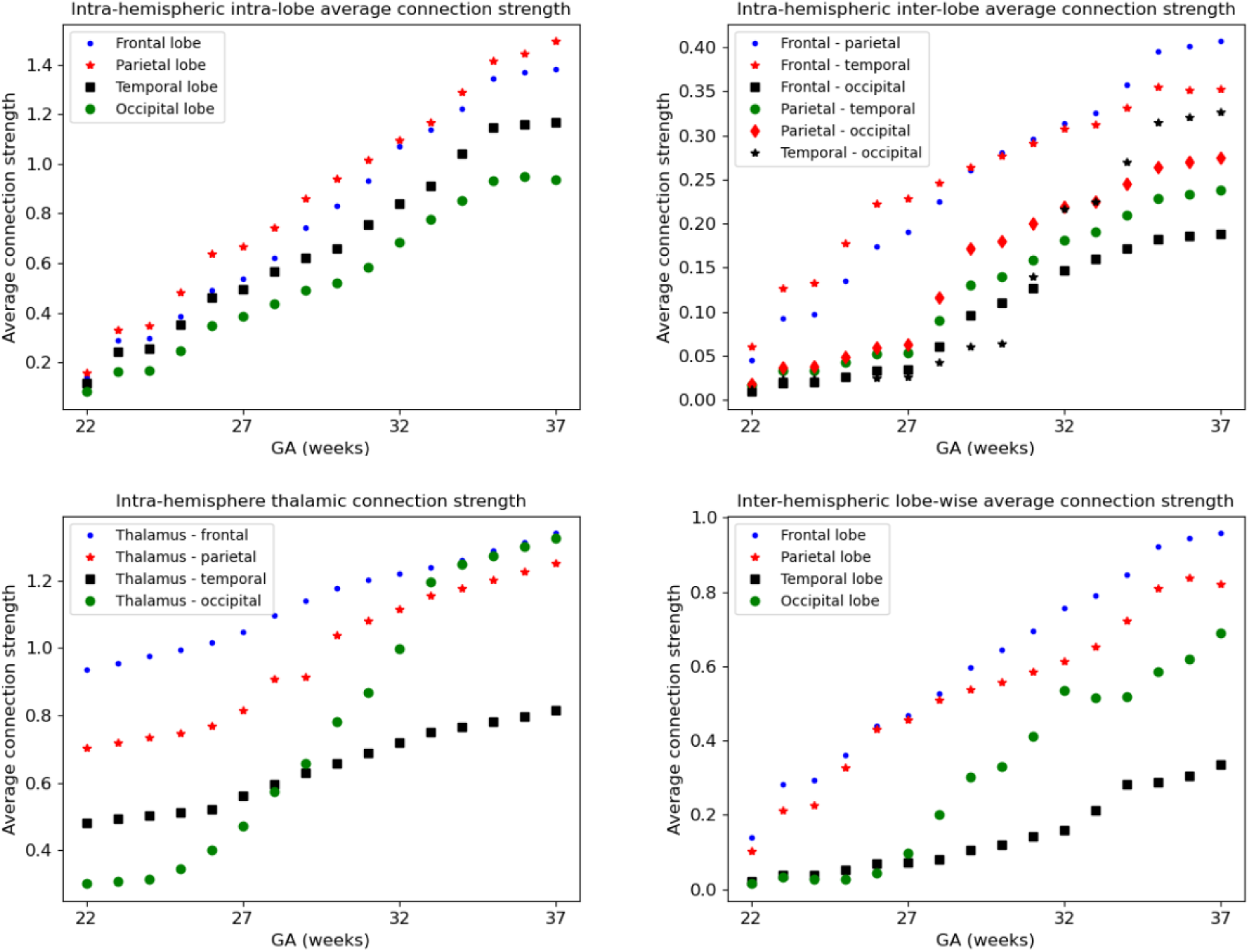
Development of structural connectivity across different brain regions computed using the connectome templates constructed with connectome aggregation method.

The developmental trajectories are largely smooth and continuous, without any abrupt transitions. Nonetheless, for interpretability, we divided this interval into five approximately equal sub-periods, each capturing key developmental transitions during this critical stage of neurodevelopment.

##### Stage 1: 22–23 gestational weeks

At this early stage, global connectivity remains weak. Subcortical–cortical connections begin to emerge, particularly those linking the thalamus and lentiform nuclei to the developing frontal and parietal cortices. The corpus callosum is present but immature. Anatomically, the genu (the anterior portion of the corpus callosum) is among the first to form, allowing interhemispheric connection between homologous frontal areas. Likewise, the body of the corpus callosum begins to connect bilateral frontal and parietal lobes. Cortico–cortical connectivity within and between the lobes remains sparse, and although tractography recovers many intra–hemispheric association streamlines, their estimated weights based on fiber bundle capacity are negligible, reflecting immature or still-developing pathways.

##### Stage 2: 24–26 gestational weeks

This stage marks an increase in the strength of association pathways. Notably, intra-hemispheric connections between frontal and parietal lobes and between frontal and temporal lobes experience strengthening. The splenium of the corpus callosum, which connects the occipital lobes across hemispheres, begins to mature during this window, though these interhemispheric occipital connections remain weaker than their frontal and parietal counterparts. Thalamocortical projections to early visual areas such as the calcarine cortex, cuneus, and lingual gyrus also emerge, marking the onset of organized input to the occipital lobe. This can be seen in the plots as the early rise in thalamus–occipital connectivity. Thalamocortical connections are of special interest during fetal development because they are among the first long-range pathways to form and play a foundational role in establishing the hierarchical organization of the cortex. These connections provide the primary sensory input to the developing brain and are thought to guide the areal differentiation of the cortex, making them critical for understanding both normative development and vulnerability to early insults.

##### Stage 3: 27–29 gestational weeks

This period is characterized by a consistent increase in the strength of nearly all connections, with several connections showing more pronounced acceleration. These include inter-hemispheric connections between the occipital lobes as well as the connections of occipital lobes to the thalami. Long-range intra-hemispheric association pathways also become more prominent, particularly those linking the occipital lobe to temporal and parietal regions. These maturing pathways form the anatomical substrate for higher cognitive and sensory integration processes that will support functions such as language, attention, and memory. Thalamocortical pathways continue to refine, particularly those targeting sensory and association cortices, contributing to the emerging coordination of large-scale brain networks.

##### Stage 4: 30–32 gestational weeks

Connectivity continues to strengthen globally, but several regional developments stand out. Interhemispheric temporal lobe connections, while still on average less strong than those of the frontal and parietal lobes, show signs of growth. Interhemispheric integration in the occipital lobe accelerates further as the splenium of the corpus callosum matures. Inter-lobe connections between the occipital and temporal lobes show a highly accelerated pace, much faster than all other intra-hemispheric inter-lobe connections at this stage, suggesting growing interdependence between visual and auditory/language processing areas. The insula, a key multisensory and salience–processing hub, exhibits increasing connectivity with multiple lobes, indicating its emerging role in integrating sensory, emotional, and cognitive information.

##### Stage 5: 33–37 gestational weeks

By late gestation, most major cortico-cortical association pathways – including those linking the temporal, occipital, and parietal lobes – are well established. Connectivity within the limbic system, particularly involving the amygdala and hippocampus, strengthens significantly, especially in their projections to the cingulate and temporal cortices. While connection strength continues to increase both within and across lobes and hemispheres, the rate of change slows down in some pathways. This may suggest that the overall scaffold of the neonatal connectome is nearing completion by term-equivalent age. Postnatal development will likely involve fine-tuning processes such as myelination, pruning, and synaptic remodeling, building upon this robust structural backbone.

The stage-wise changes observed in our data broadly align with known milestones of human fetal brain connectivity described in histological and developmental studies [121, 122, 39, 123]. During the early portion of our window (approximately 22-26 gestational weeks; Stages 1-2), the cerebral wall is dominated by the transient subplate compartment, which serves as a major site of early synaptic activity and axonal accumulation. Many long-range projections, including thalamocortical and corticocortical axons, temporarily reside or interact within this compartment before entering the cortical plate, creating a transient scaffold for early connectivity across regions [122, 39]. During this same developmental interval, thalamocortical fibers reach the subplate and accumulate there during a so-called waiting period, while the cortical plate itself remains relatively immature. These anatomical observations are consistent with the relatively weak but gradually emerging subcortical-cortical and early association connections observed in our Stages 1-2, where global connectivity remains limited but the first organized projections linking thalamic nuclei and developing cortical territories begin to appear.

From roughly the late second to early third trimester (approximately 27-37 weeks; Stages 3-5), developmental studies report a transition from this transient circuitry toward more stable cortical networks. During this interval, thalamocortical axons progressively invade the cortical plate and establish increasingly mature synaptic contacts, while corticocortical pathways and long-range association fibers expand and strengthen across the developing cortex [39, 123]. In parallel, the subplate gradually resolves as cortical circuitry becomes increasingly consolidated, and inter-hemispheric and intrahemispheric pathways, including callosal and association systems, undergo substantial growth and organization [122, 123]. This developmental progression corresponds closely with the increasingly widespread strengthening of cortico-cortical, thalamocortical, and interhemispheric connections observed in our Stages 3-5, culminating in late gestation with a relatively well-established large-scale structural axonal scaffold that precedes the extensive postnatal processes of myelination, synaptic refinement, and network specialization [123].

### 3.3 Reproducibility analysis

As described in Section 2.5, we used two different approaches to assess the reproducibility of connectome computations. First, we computed two separate connectomes, 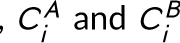, from disjoint subsets of dMRI measurements for each fetus, and computed intra-subject and inter-subject differences in terms of GED. Our analysis showed that intra-subject GEDs were significantly lower than inter-subject GEDs, with mean values of 5038±1745 and 13619±2603, respectively. For all 198 fetuses, the two connectomes derived from the independent subsets of their data were more similar to each other than to the connectomes of any other subject. This result demonstrates strong reproducibility of individual–specific structural connectivity profiles.

Secondly, we fitted linear models to graph-theoretical metrics as a function of gestational age, computed using the two measurement subsets. The resulting regression models showed no significant differences in slope or intercept across the two data halves for any metric *p* > 0.05, further confirming the robustness and reproducibility of the estimated connectomes and the derived network properties.

Our reproducibility analysis focused on consistency across independent subsets of dMRI measurements from the same fetus, demonstrating that individual-specific connectivity profiles are robust to scan-rescan variability within a fixed methodological pipeline. However, we should note that reproducibility across different computational choices, such as alternative parcellation schemes, tractography algorithms, or fiber orientation models, remains an important but separate question. Use of alternative computational methods, for example a cortical parcellation method with different definitions of regional boundaries, may also lead to variabilities in results.

### 3.4 Limitations and future work

In this study, we estimated fiber orientation distributions using single-shell CSD applied to the b= 1000 as well as DTI. We initially attempted multi-shell multi-tissue CSD (MSMT-CSD) [76], which has been successfully applied in adult and neonatal populations. However, in our fetal data, MSMT-CSD consistently failed to estimate accurate partial volume fractions for white matter, gray matter, and cerebrospinal fluid, resulting in implausibly low white matter FOD amplitudes across many regions and consequently poor tractography performance. We therefore opted for the more standard single-shell CSD and DTI approaches, which yielded visually and quantitatively superior tractography results. Nonetheless, alternative methods for fiber modeling in the fetal brain have been proposed [35, 124]. These approaches may offer advantages for specific research questions, and their systematic evaluation in the context of whole-brain structural connectome construction can be considered as a direction for future work.

Another limitation of this study is the absence of data from gestational weeks 17-21, as our earliest time point is 22 weeks. This gap reflects the limited availability of high-quality fetal dMRI data at earlier gestational ages in the dHCP dataset, due in part to greater fetal motion and smaller brain size, which also pose challenges for reliable tractography. Consequently, our findings do not capture the earliest stages of white matter pathway formation, including the initial emergence of thalamocortical and callosal fibers. Future studies targeting this earlier developmental window will be important to provide a more complete picture of fetal connectome development from mid-gestation onward.

Another consideration relates to the evaluation of our age-specific connectome templates. We benchmarked the proposed connectome aggregation method against image-space averaging by assessing how closely each template resembled individual subject connectomes of matching gestational age. While this approach is intuitive, as a template should ideally represent its target population, we acknowledge that individual fetal connectomes are inherently noisy and subject to tractography inaccuracies. The same limitations, however, apply to tractography derived from image-space-averaged data, and the question ultimately concerns which method better mitigates these imperfections to yield a faithful representation of normative development. Nevertheless, we recognize that a more definitive validation could involve comparison against independent, higher-fidelity references such as ex vivo diffusion imaging or histological data. Such validation is challenging and lies beyond the scope of the current study, but we hope to explore it in future work. Additionally, while our reproducibility analyses and concordance with histological and prior imaging findings support the biological validity of the reconstructed connectomes, we did not perform explicit validation of specific white-matter tracts against histological or ex-vivo references. Such validation, as recently demonstrated in fetal structural connectivity [125], remains an important direction for future investigation.

Furthermore, an important consideration when interpreting developmental changes in fetal dMRI connectivity is whether observed increases in connection strength reflect true biological maturation of white matter pathways (e.g., axonal packing, myelination, or synaptogenesis) or improved sensitivity of diffusion MRI to existing fibers at later gestational ages. Several factors could contribute to the latter: the smaller brain size and lower signal-to-noise ratio at earlier ages may impede tractography; incomplete myelination in the second trimester alters diffusion anisotropy and may reduce fiber orientation contrast; and fetal motion may disproportionately affect younger fetuses. Thus, while our findings of monotonic increases in network metrics are consistent with established histological timelines of axonal growth and pathway elaboration [121], we acknowledge that some portion of the observed developmental trajectory may be attributable to improved data quality and fiber resolvability with advancing gestation. Studies combining in utero dMRI with postmortem validation or advanced diffusion models that are less sensitive to myelination-dependent contrast will be important for disentangling these effects as well.

## 4 CONCLUSIONS

To the best of our knowledge, this study represents the largest investigation of the structural connectivity of fetal brain based on diffusion MRI [25]. Analysis of connectivity metrics including CPL, GE, CC, and LE revealed a monotonic increase in modular integration and segregation of the structural brain networks in this period. Our results also showed a small-world configuration, persistent across the studied period between 22 and 37 gestational weeks. Our findings showed that the connection strength for all 88 nodes considered in this analysis increased as a function of gestational age.

A technical contribution of this work was the new methods for constructing connectome templates to represent the typical development of structural networks in the fetal period. Our evaluations showed that our novel approach based on aggregation of the individual connectomes performed better than another approach based on precise spatiotemporal alignement and averaging of data in the image domain. Our analysis showed that connectome-wise averaging led to connectome templates that represented connectomes of similar-age fetuses very closely in terms of connectivity metrics and hubs. Averaging in the image space, though a sophisticated approach, resulted in connectomes that deviated from individual fetuses. This deviation is likely due to preferential weakening of certain connections and strengthening of other connections as a result of data averaging in the image space as implemented in this project. Our approach to data averaging in the image space, described in Section 2.4.2 and Figure 2, is similar to the approaches followed by many prior studies on fetal brain imaging [34, 126, 93, 127]. Therefore, our findings suggest that these widely-used approaches to constructing fetal brain templates cannot preserve the features of individual fetal brain connectome, although they may be suitable for other purposes such as studying distinct brain regions or local development of brain structure and microstructure.

Using the connectome templates generated via connectome aggregation, we characterized spatiotemporal trends in fetal brain connectivity between 22 and 37 gestational weeks. Although the developmental trajectory appeared largely continuous, we divided this period into five developmental sub-phases to highlight the key transitions in structural maturation. Around 22 weeks, thalamocortical connections are relatively much stronger than cortical-cortical connections, which are still immature both within and between hemispheres. By 37 weeks, on the other hand, dense and symmetric connections are observed within each lobe, across lobes, and between homologous regions across hemispheres. Each stage of development showed specific patterns of accelerated growth in particular connections, such as thalamocortical visual pathways, intra– and inter–hemispheric occipital–temporal associations, and insular hub emergence, reflecting the sequential establishment of large–scale functional systems during gestation.

Although the reproducibility of tractography and quantitative connectomics remains a topic of ongoing research, our reproducibility analysis yielded promising results. Connectomes reconstructed from two non-overlapping subsets of dMRI measurements for each fetus were substantially more similar to each other than to those of any other subject, demonstrating strong individual specificity. Bootstrapped assessments of nodal and edge–wise connectivity further confirmed statistically robust and reproducible patterns, as illustrated in Figure 7. Additionally, the ability of the age-specific connectome templates to predict gestational age with an average error of less than one week suggests that the structural connectome captures meaningful and temporally specific developmental signatures. Collectively, these findings indicate that, despite inherent limitations in fetal dMRI and tractography, current methodologies are capable of capturing individualized profiles of fetal brain connectivity. As such, structural connectomics may serve as a valuable framework for investigating normative and atypical patterns of prenatal brain development.

Our results also align with and extend several recent investigations of fetal brain structural connectivity. Marami et al. [69] demonstrated small-world properties in the fetal connectome using motion-robust DTI reconstruction in 21 fetuses. Calixto et al. [72] used a spatiotemporal DTI atlas to analyze fetal brain networks between 23 and 35 weeks, reporting significant weekly increases in global and local efficiency (5.83% and 5.43% per week, respectively) and the presence of small-worldness, which closely parallels our findings. The most directly comparable study is that of Chen et al. [33], who analyzed intrahemispheric corticocortical structural networks in 114 fetuses between 26 and 38 weeks using in utero dMRI. Both studies report increasing global efficiency, local efficiency, and clustering coefficient with gestational age, alongside decreasing characteristic path length and persistent small-worldness. Our study extends these findings through several methodological innovations: ensemble tractography with SIFT2 weighting, inclusion of interhemispheric connections (Chen et al. [33] excluded them due to concerns about false positives at early gestational ages), novel age-specific connectome templates, and comprehensive hub and reproducibility analyses. The physiological consistency between our results and those of Chen et al. [33], particularly regarding balanced integration and segregation of fetal brain networks, provides convergent evidence for robust developmental principles across independent cohorts and analytical pipelines. Finally, we note that Jakab et al. [29] examined network alterations in fetuses with corpus callosum agenesis compared to controls, reporting reduced clustering coefficient and altered hub centrality in pathological cases; their control group analyses provide indirect support for organized network architecture during the fetal period, though their primary focus differs from the normative developmental trajectories studied here.

Despite general agreement with existing literature, some differences between our findings and prior studies are notable. For instance, the SWI values observed in our study were lower and more stable than those reported previously, where values substantially greater than one and sharp increases with gestational age have been described [46, 128, 129, 49, 130, 51]. Similarly, although our hub analysis identified regions in agreement with previous structural and functional studies including the precuneus, thalamus, lentiform, and medial frontal cortex, we did not consistently identify the hippocampus or cingulate cortex as major hubs. These discrepancies may stem from several factors. For example, different definitions of the small–world index exist, and not all prior studies clearly report their chosen definition [86, 131, 132]. More importantly, many earlier works have used deterministic tractography and surrogate connectivity measures such as streamline count or FA [46, 49, 51, 133, 69], whereas in this study we have used anatomically constrained tractography tailored to fetal anatomy [73] and a biologically grounded measure of connection weight based on fiber bundle capacity [10, 83]. Finally, the size and quality of the data included in this study far exceed those used in prior fetal studies, allowing for more robust and generalizable conclusions. Therefore, we believe our findings offer new insights that may help reinterpret the results of earlier studies on this topic.

## 5#ACKNOWLEDGEMENTS

This research was supported in part by the National Institute of Neurological Disorders and Stroke and Eunice Kennedy Shriver National Institute of Child Health and Human Development of the National Institutes of Health (NIH) under award numbers R01HD110772, R01NS128281, R01EB036945, R01EB032708, R01EB019483, R01LM013608, R01NS10603, R01HD109395, R01EB032366 and R01EB031849. The content of this publication is solely the responsibility of the authors and does not necessarily represent the official views of the NIH. The dHCP data were provided by the developing Human Connectome Project, KCL-Imperial-Oxford Consortium funded by the European Research Council under the European Union Seventh Framework Programme (FP/2007-2013) / ERC Grant Agreement no. [319456]. We are grateful to the families who generously supported this trial.

## **6** CONFLICT OF INTEREST

The authors have no conflicts of interest.

## Notes

### Competing Interest Statement

The authors have declared no competing interest.

### Summary of Updates

New results have been added. Some old results were removed as they did not help the flow of the paper.

## references

[1] Seung S. Connectome: How the brain’s wiring makes us who we are. HMH; 2012.

[2] Sporns O, Tononi G, Kötter R. The human connectome: a structural description of the human brain. PLoS computational biology 2005;1(4):e42.

[3] Collin G, Van Den Heuvel MP. The ontogeny of the human connectome: development and dynamic changes of brain connectivity across the life span. The Neuroscientist 2013;19(6):616–628.

[4] Bullmore ET, Bassett DS. Brain graphs: graphical models of the human brain connectome. Annual review of clinical psychology 2011;7:113–140.

[5] Fornito A, Zalesky A, Breakspear M. The connectomics of brain disorders. Nature Reviews Neuroscience 2015;16(3):159–172.

[6] Shi Y, Toga AW. Connectome imaging for mapping human brain pathways. Molecular psychiatry 2017;22(9):1230–1240.

[7] Sporns O. Networks of the Brain. MIT press; 2016.

[8] Tymofiyeva O, Hess C, Xu D, Barkovich A. Structural MRI connectome in development: challenges of the changing brain. The British journal of radiology 2014;87(1039):20140086.

[9] Rossini P, Di Iorio R, Bentivoglio M, Bertini G, Ferreri F, Gerloff C, et al. Methods for analysis of brain connectivity: An IFCN-sponsored review. Clinical Neurophysiology 2019;130(10):1833–1858.

[10] Smith RE, Raffelt D, Tournier JD, Connelly A. Quantitative streamlines tractography: methods and inter-subject normalisation. Aperture Neuro 2022;p. 1–25.

[11] Zhang F, Daducci A, He Y, Schiavi S, Seguin C, Smith R, et al. Quantitative mapping of the brain’s structural connectivity using diffusion MRI tractography: a review. NeuroImage 2022;p. 118870.

[12] Sotiropoulos SN, Zalesky A. Building connectomes using diffusion MRI: why, how and but. NMR in Biomedicine 2019;32(4):e3752.

[13] Rubinov M, Sporns O. Complex network measures of brain connectivity: uses and interpretations. Neuroimage 2010;52(3):1059–1069.

[14] Assaf Y, Pasternak O. Diffusion tensor imaging (DTI)-based white matter mapping in brain research: a review. Journal of molecular neuroscience 2008;34(1):51–61.

[15] Essayed WI, Zhang F, Unadkat P, Cosgrove GR, Golby AJ, O’Donnell LJ. White matter tractography for neurosurgical planning: A topography-based review of the current state of the art. NeuroImage: Clinical 2017;15:659–672.

[16] Ciccarelli O, Catani M, Johansen-Berg H, Clark C, Thompson A. Diffusion-based tractography in neurological disorders: concepts, applications, and future developments. The Lancet Neurology 2008;7(8):715–727.

[17] Thomas C, Frank QY, Irfanoglu MO, Modi P, Saleem KS, Leopold DA, et al. Anatomical accuracy of brain connections derived from diffusion MRI tractography is inherently limited. Proceedings of the National Academy of Sciences 2014;111(46):16574–16579.

[18] Griffa A, Baumann PS, Thiran JP, Hagmann P. Structural connectomics in brain diseases. Neuroimage 2013;80:515–526.

[19] Qi S, Meesters S, Nicolay K, ter Haar Romeny BM, Ossenblok P. The influence of construction methodology on structural brain network measures: A review. Journal of neuroscience methods 2015;253:170–182.

[20] Jones DK, Knösche TR, Turner R. White matter integrity, fiber count, and other fallacies: the do’s and don’ts of diffusion MRI. Neuroimage 2013;73:239–254.

[21] Fornito A, Zalesky A, Breakspear M. Graph analysis of the human connectome: promise, progress, and pitfalls. Neuroimage 2013;80:426–444.

[22] Jones DK. Challenges and limitations of quantifying brain connectivity in vivo with diffusion MRI. Imaging in Medicine 2010;2(3):341.

[23] Drakesmith M, Caeyenberghs K, Dutt A, Lewis G, David A, Jones DK. Overcoming the effects of false positives and threshold bias in graph theoretical analyses of neuroimaging data. Neuroimage 2015;118:313–333.

[24] Yang JYM, Yeh CH, Poupon C, Calamante F. Diffusion MRI tractography for neurosurgery: the basics, current state, technical reliability and challenges. Physics in Medicine & Biology 2021;66(15):15TR01.

[25] Di Stefano M, Ciceri T, Leemans A, de Zwarte SM, De Luca A, Peruzzo D. Diffusion mri of the prenatal fetal brain: A methodological scoping review. NeuroImage 2025;p. 121453.

[26] Ouyang M, Dubois J, Yu Q, Mukherjee P, Huang H. Delineation of early brain development from fetuses to infants with diffusion MRI and beyond. Neuroimage 2019;185:836–850.

[27] Huang H, Zhang J, Wakana S, Zhang W, Ren T, Richards LJ, et al. White and gray matter development in human fetal, newborn and pediatric brains. Neuroimage 2006;33(1):27–38.

[28] Huang H, Xue R, Zhang J, Ren T, Richards LJ, Yarowsky P, et al. Anatomical characterization of human fetal brain development with diffusion tensor magnetic resonance imaging. Journal of Neuroscience 2009;29(13):4263–4273.

[29] Jakab A, Kasprian G, Schwartz E, Gruber GM, Mitter C, Prayer D, et al. Disrupted developmental organization of the structural connectome in fetuses with corpus callosum agenesis. Neuroimage 2015;111:277–288.

[30] Dubois J, Dehaene-Lambertz G, Kulikova S, Poupon C, Hüppi PS, Hertz-Pannier L. The early development of brain white matter: a review of imaging studies in fetuses, newborns and infants. Neuroscience 2014;276:48–71.

[31] Jakab A, Tuura R, Kellenberger C, Scheer I. In utero diffusion tensor imaging of the fetal brain: a reproducibility study. NeuroImage: Clinical 2017;15:601–612.

[32] Qiu A, Mori S, Miller MI. Diffusion tensor imaging for understanding brain development in early life. Annual review of psychology 2015;66:853–876.

[33] Chen R, Zhao R, Li H, Xu X, Li M, Zhao Z, et al. Development of the Fetal Brain Corticocortical Structural Network during the Second-to-Third Trimester Based on Diffusion MRI. Journal of Neuroscience 2024;44(29).

[34] Khan S, Vasung L, Marami B, Rollins CK, Afacan O, Ortinau CM, et al. Fetal brain growth portrayed by a spatiotemporal diffusion tensor MRI atlas computed from in utero images. NeuroImage 2019;185:593–608.

[35] Wilson S, Pietsch M, Cordero-Grande L, Price AN, Hutter J, Xiao J, et al. Development of human white matter pathways in utero over the second and third trimester. Proceedings of the National Academy of Sciences 2021;118(20).

[36] Pietsch M, Christiaens D, Hutter J, Cordero-Grande L, Price AN, Hughes E, et al. A framework for multi-component analysis of diffusion MRI data over the neonatal period. Neuroimage 2019;186:321–337.

[37] Hüppi PS, Dubois J. Diffusion tensor imaging of brain development. In: Seminars in Fetal and Neonatal Medicine, vol. 11 Elsevier; 2006. p. 489–497.

[38] Silbereis JC, Pochareddy S, Zhu Y, Li M, Sestan N. The cellular and molecular landscapes of the developing human central nervous system. Neuron 2016;89(2):248–268.

[39] Kostović I, Jovanov-Milošević N; Elsevier. The development of cerebral connections during the first 20–45 weeks’ gestation. Seminars in Fetal and Neonatal Medicine 2006;11(6):415–422.

[40] Bayer SA, Altman J. The human brain during the second trimester. CRC Press; 2005.

[41] Miller JA, Ding SL, Sunkin SM, Smith KA, Ng L, Szafer A, et al. Transcriptional landscape of the prenatal human brain. Nature 2014;508(7495):199–206.

[42] Innocenti GM, Price DJ. Exuberance in the development of cortical networks. Nature Reviews Neuroscience 2005;6(12):955–965.

[43] Taverna E, Götz M, Huttner WB. The cell biology of neurogenesis: toward an understanding of the development and evolution of the neocortex. Annual review of cell and developmental biology 2014;30:465–502.

[44] Yakovlev P. The myelogenetic cycles of regional maturation of the brain. Regional development of the brain in early life 1967;p. 3–70.

[45] Jakovcevski I, Filipovic R, Mo Z, Rakic S, Zecevic N. Oligodendrocyte development and the onset of myelination in the human fetal brain. Frontiers in neuroanatomy 2009;3:5.

[46] Van Den Heuvel MP, Kersbergen KJ, De Reus MA, Keunen K, Kahn RS, Groenendaal F, et al. The neonatal connectome during preterm brain development. Cerebral cortex 2015;25(9):3000–3013.

[47] Takahashi E, Folkerth RD, Galaburda AM, Grant PE. Emerging cerebral connectivity in the human fetal brain: an MR tractography study. Cerebral cortex 2012;22(2):455–464.

[48] Cao M, Huang H, He Y. Developmental connectomics from infancy through early childhood. Trends in neurosciences 2017;40(8):494–506.

[49] Song L, Mishra V, Ouyang M, Peng Q, Slinger M, Liu S, et al. Human fetal brain connectome: structural network development from middle fetal stage to birth. Frontiers in neuroscience 2017;11:561.

[50] Zhao T, Mishra V, Jeon T, Ouyang M, Peng Q, Chalak L, et al. Structural network maturation of the preterm human brain. Neuroimage 2019;185:699–710.

[51] Brown CJ, Miller SP, Booth BG, Andrews S, Chau V, Poskitt KJ, et al. Structural network analysis of brain development in young preterm neonates. Neuroimage 2014;101:667–680.

[52] Turk E, Van Den Heuvel MI, Benders MJ, De Heus R, Franx A, Manning JH, et al. Functional connectome of the fetal brain. Journal of Neuroscience 2019;39(49):9716–9724.

[53] Yap PT, Fan Y, Chen Y, Gilmore JH, Lin W, Shen D. Development trends of white matter connectivity in the first years of life. PloS one 2011;6(9):e24678.

[54] Fan Y, Shi F, Smith JK, Lin W, Gilmore JH, Shen D. Brain anatomical networks in early human brain development. Neuroimage 2011;54(3):1862–1871.

[55] Donofrio MT, Limperopoulos C, et al. Impact of congenital heart disease on fetal brain development and injury. Current opinion in pediatrics 2011;23(5):502–511.

[56] Lynch JK; Elsevier. Epidemiology and classification of perinatal stroke. Seminars in Fetal and Neonatal Medicine 2009;14(5):245–249.

[57] Linnet KM, Dalsgaard S, Obel C, Wisborg K, Henriksen TB, Rodriguez A, et al. Maternal lifestyle factors in pregnancy risk of attention deficit hyperactivity disorder and associated behaviors: review of the current evidence. American Journal of Psychiatry 2003;160(6):1028–1040.

[58] Maselko J, Sikander S, Bhalotra S, Bangash O, Ganga N, Mukherjee S, et al. Effect of an early perinatal depression intervention on long-term child development outcomes: follow-up of the Thinking Healthy Programme randomised controlled trial. The Lancet Psychiatry 2015;2(7):609–617.

[59] Sawyer A, Ayers S, Smith H. Pre-and postnatal psychological wellbeing in Africa: a systematic review. Journal of affective disorders 2010;123(1-3):17–29.

[60] Kinney DK, Munir KM, Crowley DJ, Miller AM. Prenatal stress and risk for autism. Neuroscience & Biobehavioral Reviews 2008;32(8):1519–1532.

[61] Schmithorst VJ, Votava-Smith JK, Tran N, Kim R, Lee V, Ceschin R, et al. Structural network topology correlates of microstructural brain dysmaturation in term infants with congenital heart disease. Human brain mapping 2018;39(11):4593–4610.

[62] Limperopoulos C, Tworetzky W, McElhinney DB, Newburger JW, Brown DW, Robertson Jr RL, et al. Brain volume and metabolism in fetuses with congenital heart disease: evaluation with quantitative magnetic resonance imaging and spectroscopy. Circulation 2010;121(1):26–33.

[63] Jaimes C, Rofeberg V, Stopp C, Ortinau C, Gholipour A, Friedman K, et al. Association of isolated congenital heart disease with fetal brain maturation. American Journal of Neuroradiology 2020;41(8):1525–1531.

[64] Meoded A, Poretti A, Tekes A, Flammang A, Pryde S, Huisman T. Prenatal MR diffusion tractography in a fetus with complete corpus callosum agenesis. Neuropediatrics 2011;42(03):122–123.

[65] Kasprian G, Brugger PC, Schöpf V, Mitter C, Weber M, Hainfellner JA, et al. Assessing prenatal white matter connectivity in commissural agenesis. Brain 2013;136(1):168–179.

[66] Millischer AE, Grevent D, Sonigo P, Bahi-Buisson N, Desguerre I, Mahallati H, et al. Feasibility and Added Value of Fetal DTI Tractography in the Evaluation of an Isolated Short Corpus Callosum: Preliminary Results. American Journal of Neuroradiology 2022;43(1):132–138.

[67] Huang H. Delineating neural structures of developmental human brains with diffusion tensor imaging. TheScientific-WorldJOURNAL 2010;10:135–144.

[68] Xu G, Takahashi E, Folkerth RD, Haynes RL, Volpe JJ, Grant PE, et al. Radial coherence of diffusion tractography in the cerebral white matter of the human fetus: neuroanatomic insights. Cerebral cortex 2014;24(3):579–592.

[69] Marami B, Salehi SSM, Afacan O, Scherrer B, Rollins CK, Yang E, et al. Temporal slice registration and robust diffusion-tensor reconstruction for improved fetal brain structural connectivity analysis. NeuroImage 2017;156:475–488.

[70] Calixto C, Machado-Rivas F, Karimi D, Cortes-Albornoz MC, Acosta-Buitrago LM, Gallo-Bernal S, et al. Detailed anatomic segmentations of a fetal brain diffusion tensor imaging atlas between 23 and 30 weeks of gestation. Human Brain Mapping 2023;44(4):1593–1602.

[71] Calixto C, Soldatelli MD, Li B, Vasung L, Jaimes C, Gholipour A, et al. White matter tract crossing and bottleneck regions in the fetal brain. Human brain mapping 2025;46(1):e70132.

[72] Calixto C, Machado-Rivas F, Karimi D, Velasco-Annis C, Cortes-Albornoz MC, Afacan O, et al. Population atlas analysis of emerging brain structural connections in the human fetus. Journal of Magnetic Resonance Imaging 2024;60(1):152–160.

73. [73] Calixto C, Jaimes C, Soldatelli MD, Warfield SK, Gholipour A, Karimi D. Anatomically constrained tractography of the fetal brain. arXiv preprint arXiv:240302444 2024;.

[74] Liu W, Calixto C, Warfield SK, Karimi D. Streamline tractography of the fetal brain in utero with machine learning. Imaging Neuroscience 2025;3:imag_a_00537.

[75] Christiaens D, Cordero-Grande L, Pietsch M, Hutter J, Price AN, Hughes EJ, et al. Scattered slice SHARD reconstruction for motion correction in multi-shell diffusion MRI. Neuroimage 2021;225:117437.

[76] Jeurissen B, Tournier JD, Dhollander T, Connelly A, Sijbers J. Multi-tissue constrained spherical deconvolution for improved analysis of multi-shell diffusion MRI data. NeuroImage 2014;103:411–426.

[77] Tournier JD, Calamante F, Connelly A. Robust determination of the fibre orientation distribution in diffusion MRI: non-negativity constrained super-resolved spherical deconvolution. Neuroimage 2007;35(4):1459–1472.

[78] Karimi D, Calixto C, Snoussi H, Li B, Cortes-Albornoz MC, Velasco-Annis C, et al. Detailed delineation of the fetal brain in diffusion MRI via multi-task learning. IEEE Transactions on Medical Imaging 2025;.

[79] Gholipour A, Rollins CK, Velasco-Annis C, Ouaalam A, Akhondi-Asl A, Afacan O, et al. A normative spatiotemporal MRI atlas of the fetal brain for automatic segmentation and analysis of early brain growth. Scientific reports 2017;7(1):1–13.

[80] Tournier JD, Calamante F, Connelly A, et al. Improved probabilistic streamlines tractography by 2nd order integration over fibre orientation distributions. In: Proceedings of the international society for magnetic resonance in medicine, vol. 1670 Ismrm; 2010.

[81] Aganj I, Lenglet C, Sapiro G, Yacoub E, Ugurbil K, Harel N. Reconstruction of the orientation distribution function in single-and multiple-shell q-ball imaging within constant solid angle. Magnetic resonance in medicine 2010;64(2):554–566.

[82] Descoteaux M. High angular resolution diffusion MRI: from local estimation to segmentation and tractography. PhD thesis, Université Nice Sophia Antipolis; 2008.

[83] Tahedl M, Tournier JD, Smith RE. Structural connectome construction using constrained spherical deconvolution in multi-shell diffusion-weighted magnetic resonance imaging. Nature Protocols 2025;p. 1–33.

[84] Smith RE, Tournier JD, Calamante F, Connelly A. SIFT2: Enabling dense quantitative assessment of brain white matter connectivity using streamlines tractography. Neuroimage 2015;119:338–351.

[85] Onnela JP, Saramäki J, Kertész J, Kaski K. Intensity and coherence of motifs in weighted complex networks. Physical Review E—Statistical, Nonlinear, and Soft Matter Physics 2005;71(6):065103.

[86] Humphries MD, Gurney K, Prescott TJ. The brainstem reticular formation is a small-world, not scale-free, network. Proceedings of the Royal Society B: Biological Sciences 2006;273(1585):503–511.

[87] Maslov S, Sneppen K. Specificity and stability in topology of protein networks. Science 2002;296(5569):910–913.

[88] Sanfeliu A, Fu KS. A distance measure between attributed relational graphs for pattern recognition. IEEE Transactions on Systems, Man, and Cybernetics 1983;SMC-13(3):353–362.

[89] Betzel RF, Griffa A, Hagmann P, Mišić B. Distance-dependent consensus thresholds for generating group-representative structural brain networks. Network neuroscience 2019;3(2):475–496.

[90] Betzel RF, Medaglia JD, Papadopoulos L, Baum GL, Gur R, Gur R, et al. The modular organization of human anatomical brain networks: Accounting for the cost of wiring. Network neuroscience 2017;1(1):42–68.

[91] Zhang H, Yushkevich PA, Alexander DC, Gee JC. Deformable registration of diffusion tensor MR images with explicit orientation optimization. Medical image analysis 2006;10(5):764–785.

[92] Raffelt D, Tournier JD, Fripp J, Crozier S, Connelly A, Salvado O. Symmetric diffeomorphic registration of fibre orientation distributions. Neuroimage 2011;56(3):1171–1180.

[93] Uus A, Grigorescu I, Pietsch M, Batalle D, Christiaens D, Hughes E, et al. Multi-channel 4D parametrized atlas of macro-and microstructural neonatal brain development. Frontiers in Neuroscience 2021;15:661704.

[94] Akhondi-Asl A, Warfield SK. Simultaneous truth and performance level estimation through fusion of probabilistic segmentations. IEEE transactions on medical imaging 2013;32(10):1840–1852.

95. [95] Dhollander T, Raffelt D, Connelly A. Unsupervised 3-tissue response function estimation from single-shell or multi-shell diffusion MR data without a co-registered T1 image. In: ISMRM workshop on breaking the barriers of diffusion MRI, vol. 5 Lisbon, Portugal; 2016.

[96] Tournier JD, Smith R, Raffelt D, Tabbara R, Dhollander T, Pietsch M, et al. MRtrix3: A fast, flexible and open software framework for medical image processing and visualisation. Neuroimage 2019;202:116137.

[97] Petrov D, Ivanov A, Faskowitz J, Gutman B, Moyer D, Villalon J, et al. Evaluating 35 methods to generate structural connectomes using pairwise classification. In: International Conference on medical Image Computing and Computer-Assisted Intervention Springer; 2017. p. 515–522.

[98] Page L, Brin S, Motwani R, Winograd T. The PageRank Citation Ranking: Bringing Order to the Web. Stanford InfoLab; 1999.

[99] Brandes U. On variants of shortest-path betweenness centrality and their generic computation. Social networks 2008;30(2):136–145.

[100] Bonacich P. Factoring and weighting approaches to status scores and clique identification. Journal of mathematical sociology 1972;2(1):113–120.

[101] Watts DJ, Strogatz SH. Collective dynamics of ‘small-world’networks. nature 1998;393(6684):440–442.

[102] Van Den Heuvel MP, Sporns O. Rich-club organization of the human connectome. Journal of Neuroscience 2011;31(44):15775–15786.

[103] Li L, Hu X, Preuss TM, Glasser MF, Damen FW, Qiu Y, et al. Mapping putative hubs in human, chimpanzee and rhesus macaque connectomes via diffusion tractography. Neuroimage 2013;80:462–474.

[104] Van Horn JD, Irimia A, Torgerson CM, Chambers MC, Kikinis R, Toga AW. Mapping connectivity damage in the case of Phineas Gage. PloS one 2012;7(5):e37454.

[105] Nijhuis EH, van Cappellen van Walsum AM, Norris DG. Topographic hub maps of the human structural neocortical network. PloS one 2013;8(6):e65511.

[106] Iturria-Medina Y, Sotero RC, Canales-Rodríguez EJ, Alemán-Gómez Y, Melie-García L. Studying the human brain anatomical network via diffusion-weighted MRI and Graph Theory. Neuroimage 2008;40(3):1064–1076.

[107] Gong G, He Y, Concha L, Lebel C, Gross DW, Evans AC, et al. Mapping anatomical connectivity patterns of human cerebral cortex using in vivo diffusion tensor imaging tractography. Cerebral cortex 2009;19(3):524–536.

[108] Van den Heuvel MP, Sporns O. Network hubs in the human brain. Trends in cognitive sciences 2013;17(12):683–696.

[109] van den Heuvel MI, Turk E, Manning JH, Hect J, Hernandez-Andrade E, Hassan SS, et al. Hubs in the human fetal brain network. Developmental cognitive neuroscience 2018;30:108–115.

[110] Cao M, He Y, Dai Z, Liao X, Jeon T, Ouyang M, et al. Early development of functional network segregation revealed by connectomic analysis of the preterm human brain. Cerebral cortex 2017;27(3):1949–1963.

[111] Cao M, Huang H, Peng Y, Dong Q, He Y. Toward developmental connectomics of the human brain. Frontiers in neuroanatomy 2016;10:25.

[112] Ball G, Aljabar P, Zebari S, Tusor N, Arichi T, Merchant N, et al. Rich-club organization of the newborn human brain. Proceedings of the National Academy of Sciences 2014;111(20):7456–7461.

[113] Huang H, Shu N, Mishra V, Jeon T, Chalak L, Wang ZJ, et al. Development of human brain structural networks through infancy and childhood. Cerebral Cortex 2015;25(5):1389–1404.

[114] Gao W, Zhu H, Giovanello KS, Smith JK, Shen D, Gilmore JH, et al. Evidence on the emergence of the brain’s default network from 2-week-old to 2-year-old healthy pediatric subjects. Proceedings of the National Academy of Sciences 2009;106(16):6790–6795.

[115] Gao W, Gilmore JH, Giovanello KS, Smith JK, Shen D, Zhu H, et al. Temporal and spatial evolution of brain network topology during the first two years of life. PloS one 2011;6(9):e25278.

[116] Fransson P, Åden U, Blennow M, Lagercrantz H. The functional architecture of the infant brain as revealed by resting-state fMRI. Cerebral cortex 2011;21(1):145–154.

[117] Thomason ME, Brown JA, Dassanayake MT, Shastri R, Marusak HA, Hernandez-Andrade E, et al. Intrinsic functional brain architecture derived from graph theoretical analysis in the human fetus. PLoS one 2014;9(5):e94423.

[118] Oldham S, Fornito A. The development of brain network hubs. Developmental cognitive neuroscience 2019;36:100607.

[119] Kaiser M. Mechanisms of connectome development. Trends in Cognitive Sciences 2017;21(9):703–717.

[120] Fornito A, Suarez R, Pang JC, Margulies D, van den Heuvel M, Oldham S, et al. Transmodal association hubs of the cerebral cortex: maps, models, and mechanisms. Preprint at 10.31219/osfio/tsjq6_v1 2025;.

[121] Kostović I, Sedmak G, Judaš M. Neural histology and neurogenesis of the human fetal and infant brain. Neuroimage 2019;188:743–773.

[122] Kostović I. The enigmatic fetal subplate compartment forms an early tangential cortical nexus and provides the framework for construction of cortical connectivity. Progress in neurobiology 2020;194:101883.

[123] Kostović I, Radoš M, Kostović-Srzentić M, Krsnik Ž. Fundamentals of the development of connectivity in the human fetal brain in late gestation: from 24 weeks gestational age to term. Journal of Neuropathology & Experimental Neurology 2021;80(5):393–414.

[124] Wilson S, Pietsch M, Cordero-Grande L, Christiaens D, Uus A, Karolis VR, et al. Spatiotemporal tissue maturation of thalamocortical pathways in the human fetal brain. elife 2023;12:e83727.

[125] Epihova G, Epihov DZ, Akarca D, Astle DE. The fronto-temporal cortex has increased subcortical connectivity in utero and plasticity in adulthood. The Journal of Neuroscience 2026;46(7):e0832252025.

[126] Chen R, Sun C, Liu T, Liao Y, Wang J, Sun Y, et al. Deciphering the developmental order and microstructural patterns of early white matter pathways in a diffusion MRI based fetal brain atlas. Neuroimage 2022;264:119700.

[127] Calixto C, Dorigatti Soldatelli M, Jaimes C, Pierotich L, Warfield SK, Gholipour A, et al. A detailed spatiotemporal atlas of the white matter tracts for the fetal brain. Proceedings of the National Academy of Sciences 2025;122(1):e2410341121.

[128] de Almeida JS, Meskaldji DE, Loukas S, Lordier L, Gui L, Lazeyras F, et al. Preterm birth leads to impaired rich-club organization and fronto-paralimbic/limbic structural connectivity in newborns. NeuroImage 2021;225:117440.

[129] Ratnarajah N, Rifkin-Graboi A, Fortier MV, Chong YS, Kwek K, Saw SM, et al. Structural connectivity asymmetry in the neonatal brain. Neuroimage 2013;75:187–194.

[130] Batalle D, Hughes EJ, Zhang H, Tournier JD, Tusor N, Aljabar P, et al. Early development of structural networks and the impact of prematurity on brain connectivity. Neuroimage 2017;149:379–392.

[131] Telesford QK, Joyce KE, Hayasaka S, Burdette JH, Laurienti PJ. The ubiquity of small-world networks. Brain Connec-tivity 2011;.

[132] Neal ZP. How small is it? Comparing indices of small worldliness. Network Science 2017;5(1):30–44.

[133] Shi F, Yap PT, Gao W, Lin W, Gilmore JH, Shen D. Altered structural connectivity in neonates at genetic risk for schizophrenia: a combined study using morphological and white matter networks. Neuroimage 2012;62(3):1622–1633.

